# Decomposing the sources of SARS-CoV-2 fitness variation in the United States

**DOI:** 10.1101/2020.12.14.422739

**Authors:** Lenora Kepler, Marco Hamins-Puertolas, David A. Rasmussen

## Abstract

The fitness of a pathogen is a composite phenotype determined by many different factors influencing growth rates both within and between hosts. Determining what factors shape fitness at the host population-level is especially challenging because both intrinsic factors like pathogen genetics and extrinsic factors such as host behaviour influence between-host transmission potential. These challenges have been highlighted by controversy surrounding the population-level fitness effects of mutations in the SARS-CoV-2 genome and their relative importance when compared against non-genetic factors shaping transmission dynamics. Building upon phylodynamic birth-death models, we develop a new framework to learn how hundreds of genetic and non-genetic factors have shaped the fitness of SARS-CoV-2. We estimate the fitness effects of all amino acid variants and several structural variants that have circulated in the United States between February 2020 and March 2021 from viral phylogenies. We also estimate how much fitness variation among pathogen lineages is attributable to genetic versus non-genetic factors such as spatial heterogeneity in transmission rates. Before September 2020, most fitness variation between lineages can be explained by background spatial heterogeneity in transmission rates across geographic regions. Starting in late 2020, genetic variation in fitness increased dramatically with the emergence of several new lineages including B.1.1.7, B.1.427, B.1.429 and B.1.526. Our analysis also indicates that genetic variants in less well-explored genomic regions outside of Spike may be contributing significantly to overall fitness variation in the viral population.

## Introduction

Determining what factors shape the overall fitness of a novel pathogen such as SARS-CoV-2 is a key to understanding the pathogen’s epidemiological and evolutionary dynamics. However, quantifying pathogen fitness poses a number of conceptual as well as practical challenges. The fitness of a pathogen within a host, usually defined in terms of replication or growth rates, may only have a tenuous relationship with fitness at the host population-level, which is normally defined in terms of a pathogen’s transmission potential (Handel and Rohani, 2015; Xue and Bloom, 2020). In addition to being scale-dependent, fitness is generally a composite phenotype determined by many different intrinsic (e.g. genetic) and extrinsic (e.g. environmental) factors. Several recent examples have highlighted how genetic mutations can dramatically increase the fitness of newly emerging viral pathogens including SARS-CoV, avian influenza and Ebola virus (Consortium et al., 2004; Long et al., 2016; Urbanowicz et al., 2016). At the same time, extrinsic factors such as climate and host behavior also strongly shape transmission dynamics and thereby pathogen fitness at the population-level (Shaman and Kohn, 2009; Dalziel et al., 2018; Kissler et al., 2020). Studying fitness only on one scale, or only a single component of fitness, may therefore distort our overall picture of what factors most strongly determine pathogen fitness and transmission potential.

For SARS-CoV-2, reports of novel genetic variants with enhanced infectiousness or transmissibility emerged within the first months of the global pandemic and have since received considerable attention (Korber et al., 2020a; MacLean et al., 2020b; Tang et al., 2020). Early on, the most notable of these variants was the D614G mutation in the receptor binding domain of the Spike glycoprotein that binds human ACE2 receptors during cell entry. This variant spread rapidly around the globe in the spring of 2020 and apparently out-competed other viral genotypes that were already established in several locations (Korber et al., 2020b). While experimental studies showed that the D614G variant increases cellular infectivity and viral replication rates (Korber et al., 2020b; Plante et al., 2020; Zhang et al., 2020), its impact on population-level fitness remains less clear with fitness estimates ranging from low to moderately large benefits (Leung et al., 2020a; Volz et al., 2020).

In late 2020, several new variants of SARS-CoV-2 with increased transmissibility and potential antigenic escape mutations emerged, including lineage B.1.1.7 in the UK (Volz et al., 2021; Davies et al., 2021), B.1.351 in South Africa (Tegally et al., 2020) and P.1 in Brazil (Naveca et al., 2021). All of these emerging variants were subsequently introduced into the United States as early as October or November, 2020 (Larsen and Worobey, 2021; Washington et al., 2021). However, quantifying the fitness of these lineages in the US and their impact on national-level epidemic dynamics poses a considerable challenge due to the rapidly evolving epidemic landscape in the US. In addition to introduced variants, new “domestic” variants have emerged such as B.1.427/B.1.429 in California and B.1.526 in New York (Deng et al., 2021; Walensky et al., 2021; Zhang et al., 2021). At the same time, older lineages like B.1.2 remain dominant across large geographic regions and have as yet not been replaced by newer variants (Pater et al., 2021). Furthermore, multiple lineages have independently acquired amino acid mutations suspected to increase transmission potential or escape immunity, including Spike E484K, Spike N501Y and Spike Q677P/H (Hodcroft et al., 2021; Martin et al., 2021), suggesting that lineages are adapting through convergent evolution.

While the fitness effect of genetic variants can be precisely quantified within hosts in controlled lab experiments (Urbanowicz et al., 2016; Muth et al., 2018; Zhang et al., 2020), laboratory conditions may not faithfully mimic host environments and immune responses encountered during natural infections. Moreover, due the scale-dependence of fitness, increased cellular infectivity or replication rates may not scale up to increase transmission potential between hosts, especially if within-host growth rates already produce sufficient viral loads or optimize a tradeoff between virulence and transmission (Fraser et al., 2007; Alizon et al., 2009; Ke et al., 2020). Thus, in order to provide a definitive answer about the epidemiological significance of a novel pathogen variant, fitness also needs to be quantified at the between-host or population-level.

Fitness at the population level can be inferred based on the evolutionary dynamics of pathogen variants in the host population. For example, the growth rate of alternate variants can be estimated from time series of variant frequencies or pathogen phylogenies as a surrogate for fitness (Foll et al., 2015; Kühnert et al., 2018). However, because fitness is a composite phenotype determined by multiple factors, inferring the fitness effect of a single feature such as a mutation can be easily confounded by other factors shaping pathogen fitness if these confounding factors are not accounted for. For example, a mutation of interest may be linked to other non-neutral mutations in the same genetic background and thereby confound estimates of the mutation’s fitness effect by altering the background fitness of pathogen lineages carrying the mutation (Illingworth and Mustonen, 2012; Neher, 2013). Extrinsic factors such as climate and host behavior also strongly shape transmission dynamics (Dalziel et al., 2018; Kissler et al., 2020), such that a novel variant may increase rapidly in frequency and appear to have a fitness advantage simply by being in the right host population at the right time.

Viral phylogenies offer a promising way to estimate pathogen fitness while controlling for multiple confounding factors. On average, a pathogen lineage with increased population-level fitness will be transmitted more frequently and have a higher probability of persisting through time. More fit lineages will therefore have a higher branching rate in the phylogeny and leave behind more sampled descendants. The fitness of a viral lineage can therefore be inferred from its branching pattern in a phylogeny using phylodynamic approaches such as birth-death models (Neher et al., 2014). Multi-type birth-death (MTBD) models extend this basic idea by allowing the birth and death rate of lineages, and thereby fitness, to depend on a lineage’s state or type, which may represent its genotype or any other *feature* representing a discrete character trait (Maddison et al., 2007; Stadler and Bonhoeffer, 2013; Kühnert et al., 2018). Here we develop a phylodynamic inference framework that builds on earlier MTBD models to allow the fitness of a lineage to depend on multiple evolving traits or features (Rasmussen and Stadler, 2019). In this framework, we first reconstruct ancestral states for all features that potentially predict fitness and then use a *fitness mapping function* to translate a lineage’s reconstructed ancestral features into its expected fitness. We also develop a new approach that combines recent advances in machine learning with likelihood-based statistical inference under a birth-death model to learn this fitness mapping function from a phylogeny with reconstructed ancestral features.

We apply this new phylodynamic framework to learn what genetic as well as extrinsic features determine the fitness of SARS-CoV-2 at the host population-level in the United States. This approach allows us to estimate the fitness effects of a large number of genetic variants while accounting for confounding factors such as background spatial heterogeneity in transmission. This approach also allows us to explore the relative importance of different features to overall pathogen fitness by decomposing or partitioning fitness variation among lineages into parts attributable to different components of fitness. We therefore obtain a clearer picture of what factors have most strongly shaped the fitness of SARS-CoV-2 lineages circulating in the US.

## Results

### Phylogenetic and ancestral state reconstruction

We originally analyzed a data set containing 22,416 SARS-CoV-2 whole genome sequences sampled in the United States prior to September 1st, 2020 (Pre-2020-09 data). Since the evolutionary dynamics of SARS-CoV-2 underwent a dramatic transition in late 2020, we subsequently performed an updated analysis on an additional 66,339 sequences sampled in the US between Sept 1st, 2020 and March 1st, 2021 (Post-2020-09 data). We combine these two data sets (Combined data) for some analyses below. Dated or time-calibrated maximum likelihood phylogenetic trees were reconstructed from whole genome sequences in each data set. For all sampled viruses, we also assembled a set of features that potentially predict fitness, including both genetic and non-genetic, environmental features. The genetic features include amino acid variants (AAVs) in coding regions spanning the SARS-CoV-2 genome as well as structural (deletion) variants. The non-genetic features include each sample’s spatial location both at the level of US state and geographic region as determined by the US Department of Health and Human Services. Ancestral states for all features were then reconstructed for each node in the ML phylogeny. Thus, for each lineage in the phylogeny we obtain a vector of categorical variables representing ancestral features which we use to predict a lineage’s fitness.

### Background sampling and transmission heterogeneity

Because phylodynamic estimates will inevitably depend on what pathogens are sampled for genomic sequencing, we first estimated how sampling efforts varied across the US by time and geographic region. Sampling fractions were estimated based on the number of whole-genome sequences submitted to GISAID relative to the total number of COVID infections imputed based on reported COVID deaths (see Methods). Overall, sampling fractions were extremely variable over the first eight months of the pandemic, but have become less variable and increased steadily over time since fall 2020 (Figure 1A). When averaged across all times and regions, the mean sampling fraction is estimated to be 0.14%.

**Figure 1.**
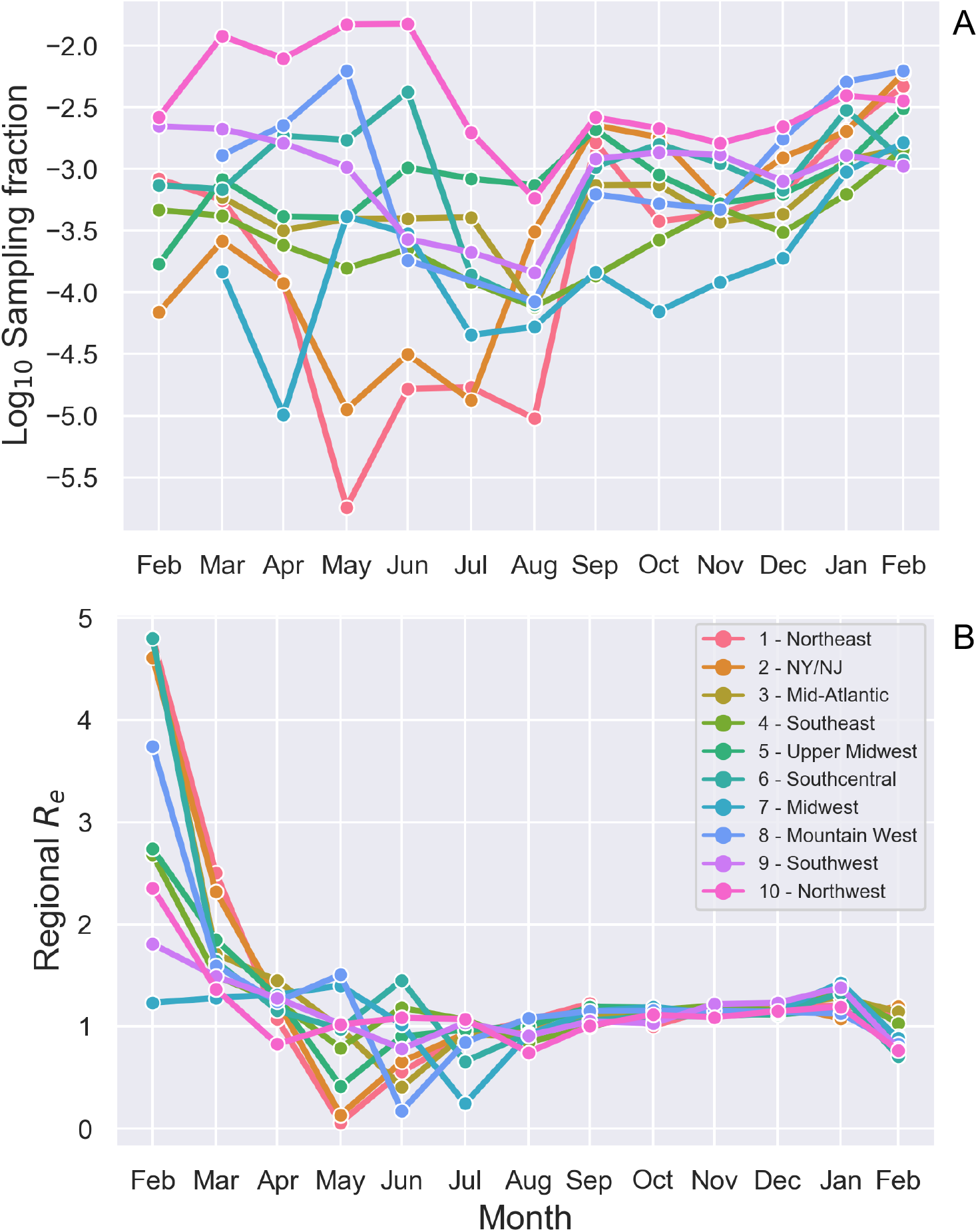
Background spatiotemporal heterogeneity in sampling fractions and effective reproductive number *R*_*e*_ of SARS-CoV-2 in the US. (A) Sampling fractions estimated based on the number of full viral genomes deposited to GISAID relative to the estimated number of total COVID infections in each region and time interval. (B) Effective reproductive number *R*_*e*_ estimates from the ML SARS-CoV-2 phylogeny. A regional transmission effect was estimated for each region and time interval, which was then used to rescale the estimated base transmission rate to compute *R*_*e*_. The base transmission rate was estimated to be 0.184 per day, which assuming a constant recovery/removal rate of 0.14 per day yields an estimate of the basic reproduction number *R*_0_ = 1.31. States are grouped into the geographic regions designated by the US Department of Health and Human Services.

Before considering models that include genetic variants as fitness-predicting features, we considered several models accounting for background spatial and temporal variability in transmission, which could otherwise confound fitness estimates. The best fitting model allowed transmission rates to vary by both monthly time interval and geographic region (see Model Selection). We therefore use a model that directly accounts for time-varying regional transmission rates and time-varying regional sampling fractions (as estimated in Figure 1A) in all subsequent analysis.

Using our phylodynamic birth-death model, we estimated how background transmission rates varied across geographic regions from the SARS-CoV-2 phylogeny. Figure 1B illustrates the changing transmission dynamics in terms of the effective reproductive number *R*_*e*_ for each region. Estimated transmission rates and *R*_*e*_ peak in February 2020, substantially earlier than peaks in reported cases. This pattern has been reported in other phylodynamic studies (Fauver et al., 2020; Nadeau et al., 2020; Ragonnet-Cronin et al., 2020), and may reflect considerable undetected transmission as well as lags in reporting before routine testing began. Transmission rates then remain low through the spring and early summer 2020 but are extremely variable across regions, likely reflecting the extreme variability in imputed sampling fractions during this same time period. Transmissions rates then steadily increase through late summer and fall of 2020 before declining in early 2021, consistent with trends observed in case report data.

### Fitness effects of genetic variants

We next estimated the fitness effect of genetic variants while controlling for background heterogeneity in transmission rates and sampling fractions. We consider the fitness effect of 66 amino acid variants (AAVs) in the Pre-2020-09 data and 110 AAVs in the Post-2020-09 data. However, in both data sets several variants are tightly linked and nearly always co-occur together (Supp. Figure 1), leading to strong collinearity among features in our model. We therefore encode sets of linked variants with correlation coefficients greater than 0.95 as single features.

Fitness effects were estimated under a model where each variant has a multiplicative effect on the base transmission rate of a lineage such that a neutral variant has a fitness effect of 1.0 and deleterious or beneficial mutants have fitness effects less than or greater than 1.0, respectively. These fitness effects therefore also directly quantify the variant’s effect on the *R*_*e*_ of pathogen lineages with the variant. We only consider a variant to be significantly deleterious or beneficial if the estimated 95% credible interval (CI) does not overlap with 1.0.

For the Pre-2020-09 data, most amino acid variants are inferred to be neutral, with maximum likelihood estimates (MLE) of fitness effects close to 1.0 and 95% CIs overlapping 1.0 (Figure 2). Estimated fitness effects are generally consistent across 10 bootstrapped phylogeny replicates. Variants with larger positive fitness effects (*>*1.05) are generally rare mutations or have wide confidence intervals surrounding the MLE. AAVs with large and significant positive effects are summarized in Table 1.

**Table 1.** Amino acid variants with significantly positive fitness effects in the Pre-2020-09 data.

| Variant | MLE | 95% CI | Frequency |
| --- | --- | --- | --- |
| nsp2 A336V | 1.15 | 1.13-1.22 | 0.006 |
| nsp2 G199E | 1.223 | 1.13-1.26 | 0.005 |
| nsp3 K384N | 1.068 | 1.06-1.12 | 0.015 |
| nsp13 P47L | 1.113 | 1.01-1.13 | 0.006 |
| S Q675R | 1.17 | 1.14-1.22 | 0.005 |
| ORF9 S194L | 1.13 | 1.01-1.24 | 0.048 |
| ORF9 T205I | 1.177 | 1.05-1.24 | 0.007 |

**Figure 2.**
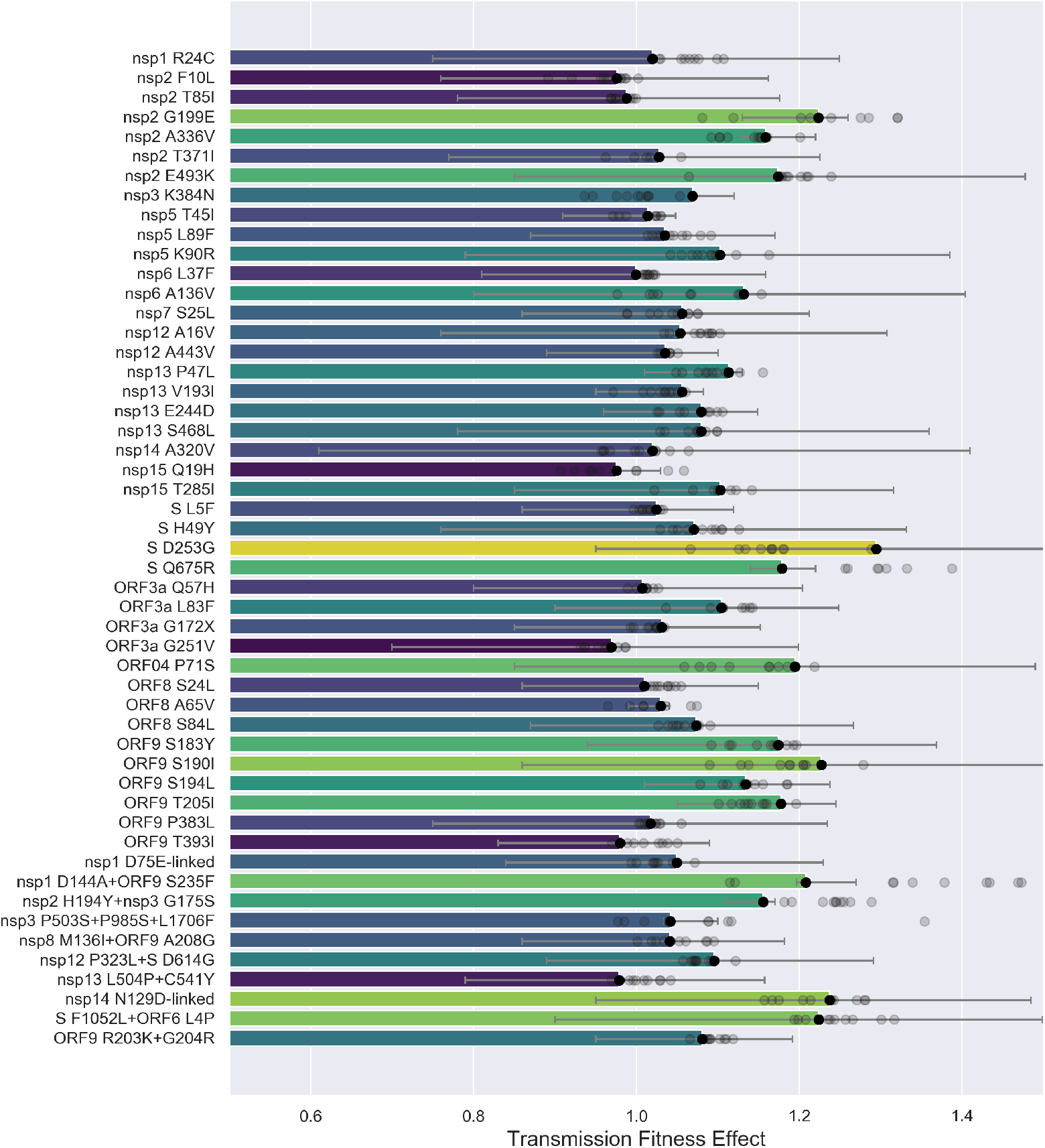
Estimated transmission fitness effects of amino acid variants in the Pre-2020-09 data. Fitness effects are jointly estimated under a model of multiplicative fitness, such that neutral variants have a fitness of one. Variants are ordered from top to bottom by their genomic position. Bars are colored according to the maximum likelihood estimate (MLE) of each variant’s fitness effect. Capped lines indicate the 95% credible interval around the MLE. The MLE of each fitness effect is also shown for 10 replicate bootstrap trees as transparent circles. Sets of strongly linked variants are grouped together as single features to avoid collinearity among features. The nsp1 D75E-linked set includes nsp1 D75E, nsp3 P153L, nsp14 F233L and ORF8 V62L; the nsp14 N129D-linked set includes nsp14 N129D, nsp16 R216C, ORF3a G172V, ORF9 P199L and ORF9 P67S. The nsp14 N129D-linked set is referred to as the B.1.2 linked set below.

The Spike D614G variant nearly always co-occurs with the P323L variant in nsp12 (RdRp), so we consider these two variants together as a single feature. Despite rapidly increasing in frequency in the spring of 2020 (Figure 4A), the Spike D614G + nsp12 P323L variant is estimated to have only a modest fitness benefit of 1.095 with a fairly wide 95% CI of 0.89-1.29. Simulations using a two-strain epidemiological model show that a transmission fitness effect of this magnitude is insufficient to explain D614G’s rapid increase in frequency during the spring of 2020. Even if D614G entered the US through external introductions at a much higher rate than the ancestral 614D variant, D614G would have required a fitness advantage much larger than 10% to rise so rapidly in frequency (Supp. Figure 3). We therefore explore other plausible explanations for D614G’s rapid rise below.

For the Post-2020-09 data, most individual AAVs are again estimated to be approximately neutral (Figure 3). Only one AAV, Spike A701V, is estimated to be significantly deleterious with a fitness effect of 0.937 (95% CI: 0.91-0.98). However, there are a relatively large number of AAVs with significant positive fitness effects between 1.05 and 1.10, especially in nsp3, ORF3a and ORF9 (Table 2).

**Figure 3.**
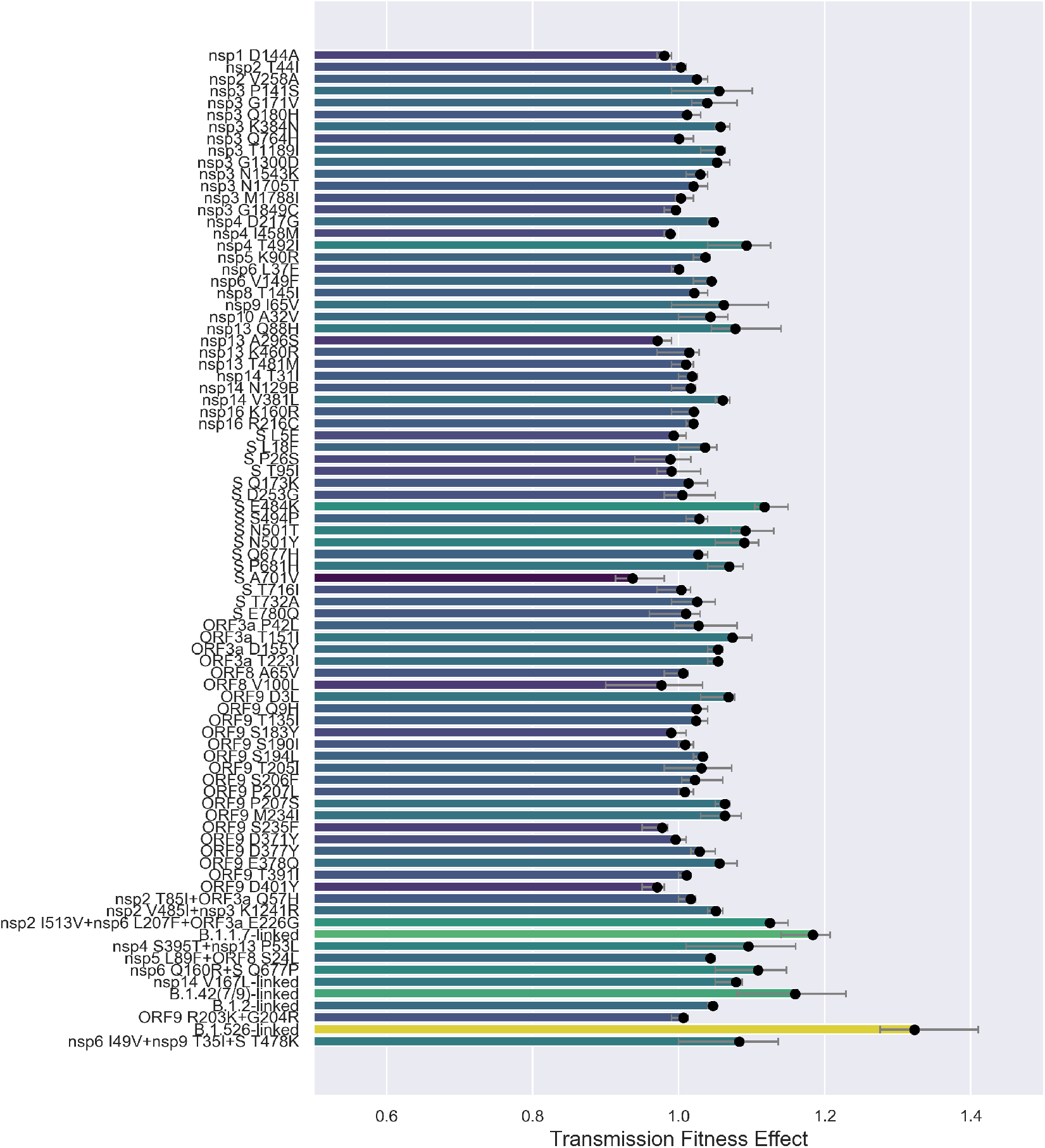
Estimated transmission fitness effects of amino acid variants in the Post-2020-09 data. The fitness of each AAV is reported as a multiplicative effect on the base transmission rate. The B.1.1.7-linked set includes nsp3 T183I, nsp3 A890D, nsp3 I1412T, S A570D, S D1118H, S S982A, ORF8 Q27, ORF8 R52I, ORF8 Y73C. The nsp14 V167L-linked set includes nsp12 V776L, nsp14 V167L, ORF3a S180P and ORF9 Q389L. The B.1.526-linked set includes nsp13 D260Y, S S13I, S W152C and S L452R. The B.1.2-linked set includes nsp14 N129D, ORF3a G172V, ORF9 P67S and ORF9 P199L.

**Table 2.** Amino acid variants outside of Spike with significantly positive fitness effects in the Post-2020-09 data

| Variant | MLE | 95% CI | Frequency |
| --- | --- | --- | --- |
| nsp3 K384N | 1.057 | 1.05-1.07 | 0.016 |
| nsp3 T1189I | 1.057 | 1.03-1.06 | 0.012 |
| nsp3 G1300D | 1.052 | 1.05-1.07 | 0.023 |
| nsp4 T429I | 1.093 | 1.04-1.13 | 0.019 |
| nsp13 Q88H | 1.077 | 1.04-1.14 | 0.012 |
| nsp14 V381L | 1.06 | 1.05-1.07 | 0.014 |
| ORF3a T151I | 1.074 | 1.07-1.11 | 0.015 |
| ORF3a D155Y | 1.053 | 1.04-1.06 | 0.018 |
| ORF3a T223I | 1.053 | 1.04-1.06 | 0.033 |
| ORF3a E226G | 1.124 | 1.12-1.15 | 0.014 |
| ORF9 D3L | 1.068 | 1.03-1.08 | 0.015 |
| ORF9 P207S | 1.063 | 1.05-1.07 | 0.015 |
| ORF9 M234I | 1.063 | 1.03-1.086 | 0.054 |
| ORF9 E378Q | 1.055 | 1.05-1.08 | 0.012 |

**Table 3.**
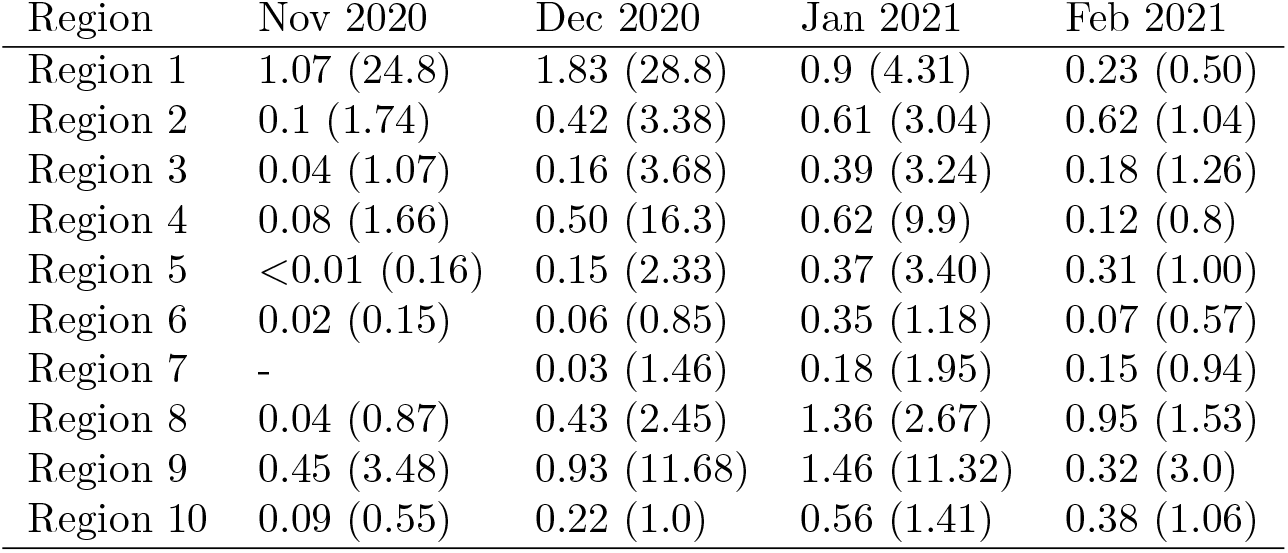
SGTF-specific sampling fractions (as percentages) and the ratio of SGTF versus non-SGTF sampling fractions (parentheses)

| Region | Nov 2020 | Dec 2020 | Jan 2021 | Feb 2021 |
| --- | --- | --- | --- | --- |
| Region 1 | 1.07 (24.8) | 1.83 (28.8) | 0.9 (4.31) | 0.23 (0.50) |
| Region 2 | 0.1 (1.74) | 0.42 (3.38) | 0.61 (3.04) | 0.62 (1.04) |
| Region 3 | 0.04 (1.07) | 0.16 (3.68) | 0.39 (3.24) | 0.18 (1.26) |
| Region 4 | 0.08 (1.66) | 0.50 (16.3) | 0.62 (9.9) | 0.12 (0.8) |
| Region 5 | <0.01 (0.16) | 0.15 (2.33) | 0.37 (3.40) | 0.31 (1.00) |
| Region 6 | 0.02 (0.15) | 0.06 (0.85) | 0.35 (1.18) | 0.07 (0.57) |
| Region 7 | - | 0.03 (1.46) | 0.18 (1.95) | 0.15 (0.94) |
| Region 8 | 0.04 (0.87) | 0.43 (2.45) | 1.36 (2.67) | 0.95 (1.53) |
| Region 9 | 0.45 (3.48) | 0.93 (11.68) | 1.46 (11.32) | 0.32 (3.0) |
| Region 10 | 0.09 (0.55) | 0.22 (1.0) | 0.56 (1.41) | 0.38 (1.06) |

Within the Spike domain, the putative antigenic escape mutation S E484K is estimated to have a significant fitness advantage (MLE: 1.117; 95% CI: 1.10-1.15). Two receptor binding domain mutations at position 501 with increased ACE2 binding avidity are also estimated to have significant positive fitness effects: S N501T (MLE: 1.091; 95% CI: 1.07-1.13) and S N501Y (MLE: 1.090; 95% CI: 1.05-1.11). The S Q677P variant, which has arisen in multiple genetic backgrounds (Pater et al., 2021; Hodcroft et al., 2021), linked with nsp6 Q160R is likewise estimated to have a significant fitness advantage (MLE: 1.109; 95% CI: 1.05-1.15), while the related mutation S Q677H is estimated to have a much smaller advantage (MLE: 1.026; 95% CI: 1.02-1.04). Finally, S P681H which has arisen multiple times including in B.1.1.7 and may aid cell entry by increasing the efficiency of furin cleavage (Garry et al., 2021), is estimated have a fitness effect of 1.069 (95% CI: 1.04-1.08).

Overall, the genetic features with the largest positive fitness effects in the Post-2020-09 data are all sets of linked AAVs associated with major lineages. The B.1.526-linked variants are estimated to have the largest fitness effect (MLE: 1.322; 95% CI: 1.27-1.41), followed by the set of the nine B.1.1.7-linked variants (MLE: 1.183: 95% CI: 1.14-1.21) and then the B.1.427 and B.1.429-linked variants including Spike L452R (MLE: 1.159; 95% 1.08-1.229). Unfortunately, due to tight genetic linkage among these mutations, we are unable to determine whether individual amino acid variants contribute disproportionately to the fitness of these lineages.

### Explaining the dominance of the Spike D614G variant

If the Spike D614G variant is not itself strongly beneficial as our fitness estimates suggest, what explains the rapid increase in the frequency of the D614G variant across the US? Stochastic processes including founder effects alone seem implausible given that the 614G variant appears to have out-competed and replaced the ancestral 614D variant even in geographic locations where the 614G variant arrived after the 614D variant (Korber et al., 2020b). We therefore consider two alternative hypotheses for the success of 614G: (1) the 614G variant gained an advantage by occurring in genetic backgrounds with higher fitness on average than the 614D variant; or (2) the 614G variant tended to occur in geographic locations with higher transmission rates on average.

We estimated the average background fitness of lineages with either the 614D or 614G variant, discounting the fitness effects of the Spike 614 variants themselves. Lineages with the 614G variant have an average background fitness that is 10.6% higher than the 614D variant. After partitioning total background fitness into genetic and spatial components, the 614G variant occurs in genetic backgrounds with 7.1% higher fitness. The genetic background fitness advantage of 614G lineages derives mostly from the ORF8 S84L variant, which we estimate had a fitness advantage of 1.073 but with a high degree of uncertainty (95% CI: 0.87-1.26). However, the ORF8 S84L variant almost always occurs in the same genetic background as D614G, so it is unclear whether this fitness advantage should be attributed exclusively to S84L or to the overall genetic fitness background of lineages with 614G

In addition to its genetic background, the 614G variant occurs in spatial backgrounds (i.e. geographic regions) with 3.4% higher fitness on average, although this average conceals the fact that the spatial background fitness advantage of the 614G variant was initially more than 10% due to being in geographic regions with higher average transmission rates during the earliest stages of the pandemic, but this spatial advantage dissipated over time (Figure 4). Directly comparing phylogenies with reconstructed ancestral states for the 614 variants with ancestral geographic locations makes clear that lineages carrying the 614G variant tended to be in locations like Region 2 (NY and NJ) and Region 5 (upper Midwest) with the highest transmission rates during the earliest stages of the pandemic (Supp. Figure 4)..

**Figure 4.**
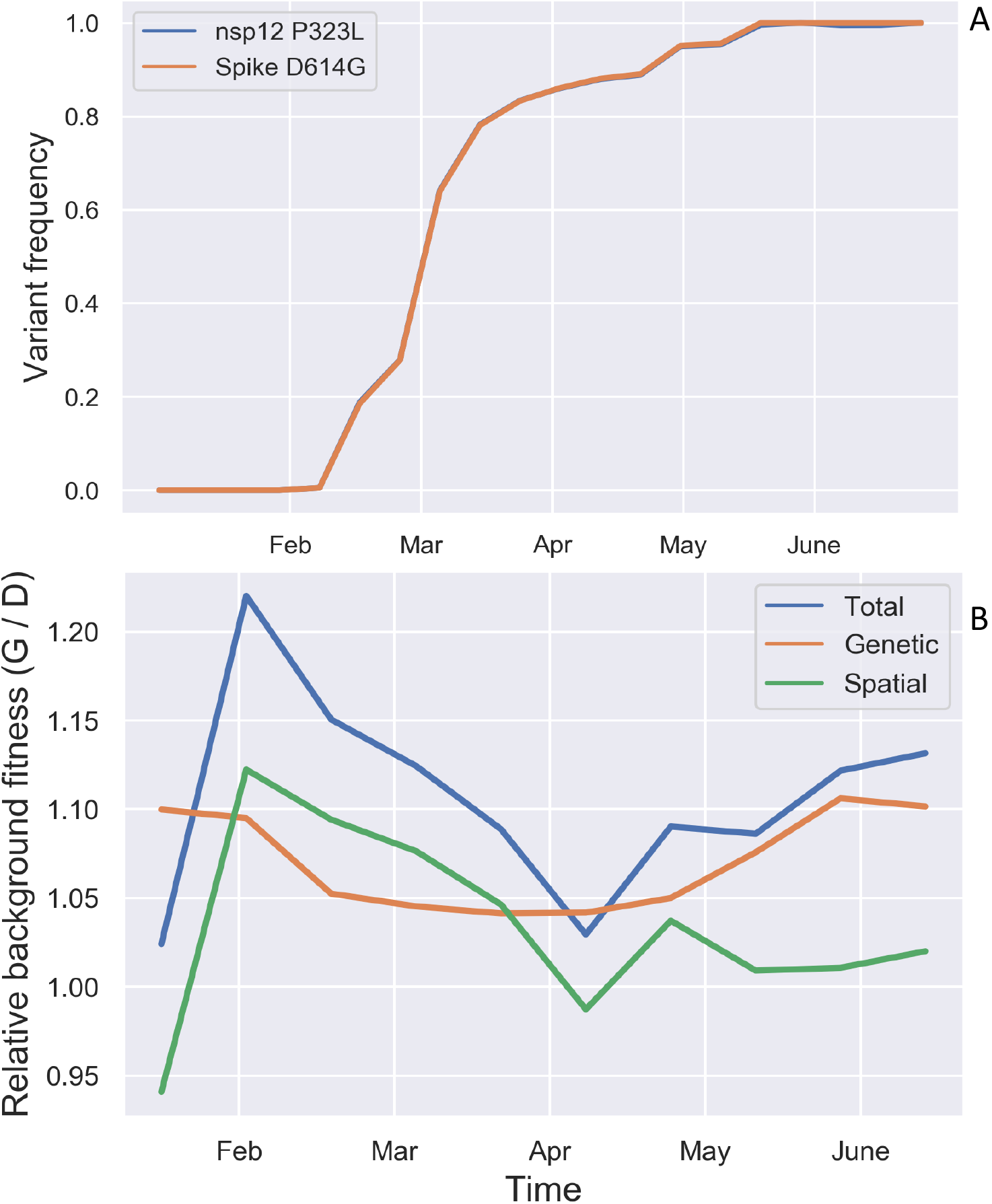
Evolutionary dynamics and background fitness of the Spike 614 variants. **(A)** Frequency of lineages carrying the Spike D614G and nsp12 P323L variants over time relative to all lineages in the ML phylogeny. These two variants are tightly linked so that they largely share the same evolutionary trajectory. **(B)** Relative background fitness of lineages with the Spike 614G variant versus the 614D variant. Background fitness was computed by averaging the fitness of all lineages with either variant present in the ML phylogeny at each time point. Total background fitness was then further split into a spatial and genetic component. Relative fitness is only shown up to July 1st, 2020 as the 614D variant was not sampled after this date.

The above analysis suggests that while lineages carrying the 614G variant may have had a small genetic fitness advantage, the 614G variant’s rapid rise in frequency across the US was largely driven by establishing first in regions with higher average transmission rates. This can be seen by comparing the cumulative number of branching events in the phylogeny for lineages with the 614D or 614G variant. Using branching events as a proxy for transmission events, lineages with the 614G variant branch more often first in Region 2 and then subsequently in all other regions (Supp. Figure 5). Nevertheless, this pattern alone does not necessarily imply that the 614G variant has an intrinsic fitness advantage or elevated transmission rate as the 614G variant is also imported more frequently into each region than the D variant (Supp. Figure 6). To place the variants on more equitable footing, we therefore compare the branching/transmission rate of the variants *per lineage*, which accounts for the the fact that the total number of lineages with the 614G variant in a given region may be higher due to either a higher transmission rate or importation rate. Contextualizing variant dynamics in this way, it becomes very clear that neither variant has a consistently higher branching rate through time (Figure 5), supporting our model-based inference that the 614G variant may not have a major intrinsic fitness advantage. Averaging over all regions and time intervals up to May 1st, after which the 614D variant is rarely sampled, the branching rate of the 614G variant (mean = 0.13 per week) is 13% higher than the 614D variant (mean = 0.115 per week), consistent with our model-based estimate of a ∼10% fitness advantage, but these means are not significantly different (Welch’s t-test = -1.42; p-value = 0.15).

**Figure 5.**
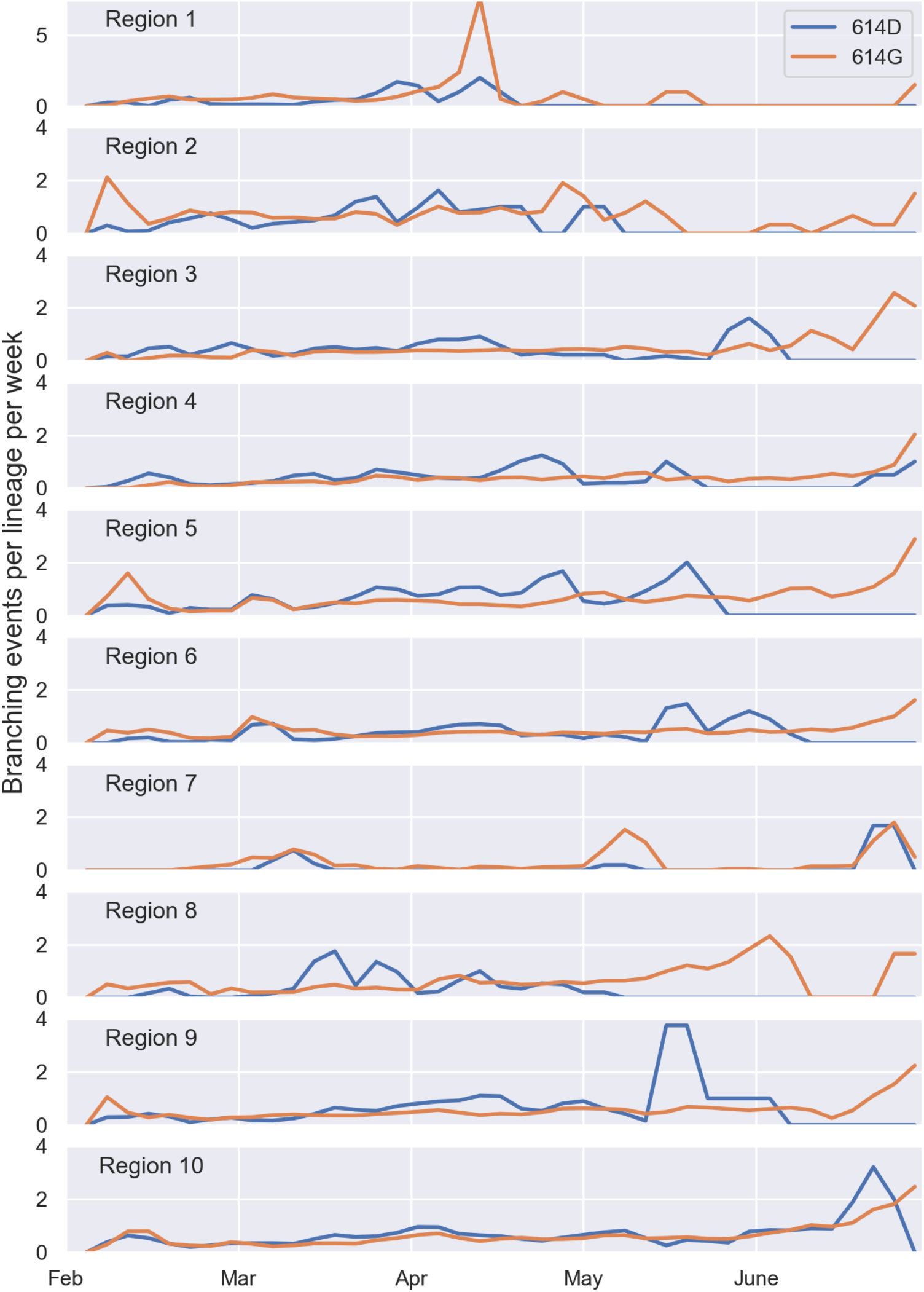
Branching rates per lineage in each region for lineages with the Spike 614D versus 614G variant. Branching rates are reported here as the number of branching events per week in the ML phylogeny for lineages with either variant.

### Fitness of major lineages circulating in the US

Because the genetic features with the largest fitness effects are all linked sets of AAVs associated with emerging lineages, we also estimated the fitness of major lineages circulating in the US as of March, 2021. Assignment of viruses to lineages is based on the Pangolin (PANGO) naming scheme (Rambaut et al., 2020). For our purposes, we only consider lineages with at least 1,000 samples in the Post-2020-09 data, plus B.1.526 (n=797). Note, that this excludes some variants of concern that are still rare in the US, including the “South African” variant B.1.351 and the “Brazilian” variant P.1.

Similar to the AAVs considered above, we estimated a multiplicative fitness effect on the base transmission rate for each lineage. The estimated fitness of each lineage is shown in Figure 6A. Relative to B.1, several common lineages are estimated to have moderate transmission fitness advantages, including B.1.2 (MLE: 1.085; 95% CI: 1.08-1.10), B.1.222 (MLE: 1.097; 95% CI: 1.05-1.12), B.1.234 (MLE: 1.076; 95% CI: 1.07-1.09) and B.1.243 (MLE: 1.041; 95% CI: 1.03-1.05).

**Figure 6.**
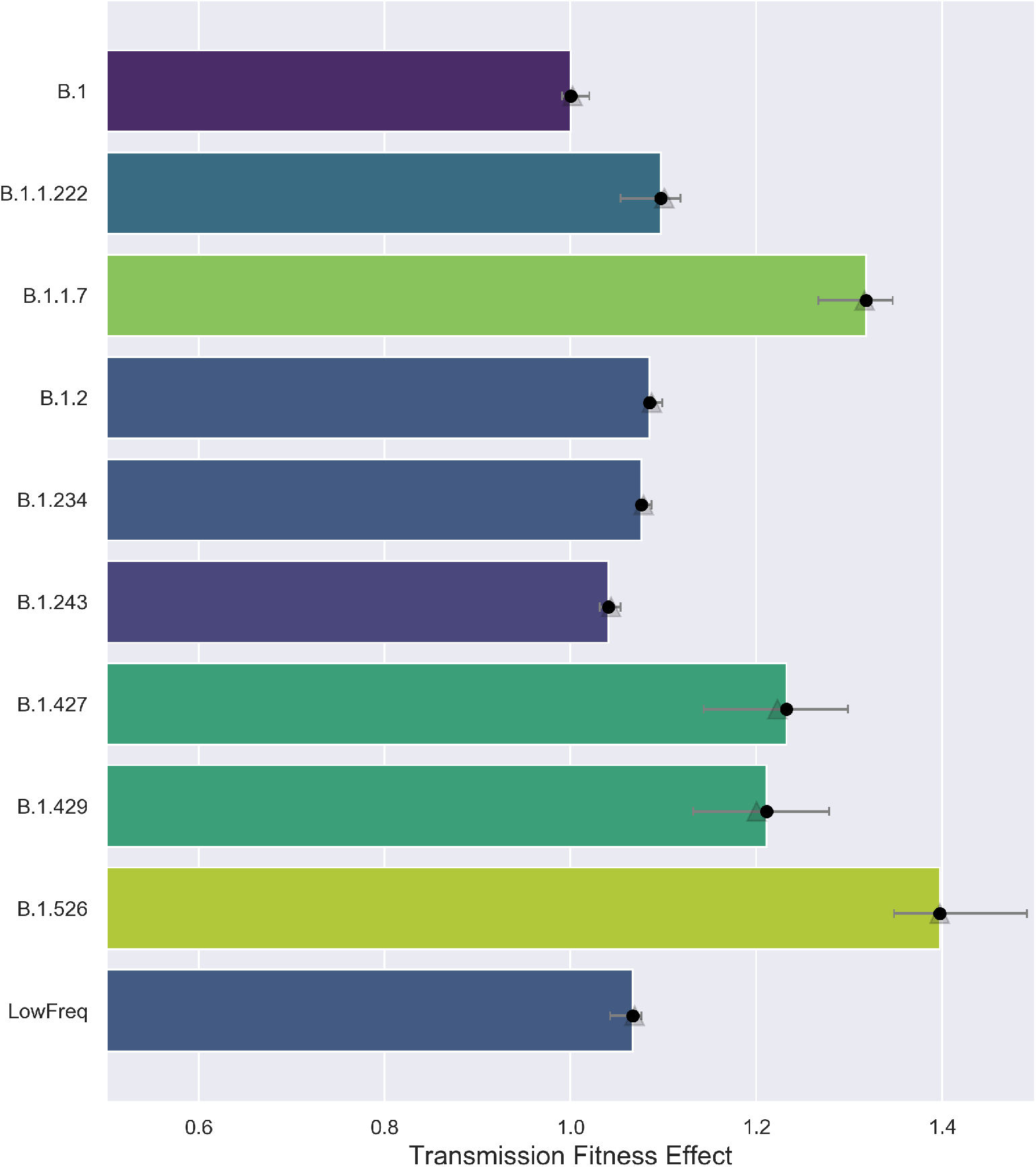
Estimated fitness of major lineages circulating in the US. The fitness of each lineage is reported as a multiplicative effect on the base transmission rate shared among all lineages. Fitness effects were estimated before (solid circles) and after (shaded triangles) accounting for SGTF-variant sampling biases by allowing lineages with the ΔH69/V70 deletion mutation to have their own SGTF-variant specific sampling fraction.

Four lineages are estimated to have a significant transmission fitness advantage. The order of these lineages’ fitness follows the same rank order as their associated sets of linked AAVs. B.1.427 and B.1.429, two sister lineages that first appeared in California and carry the Spike L452R mutation (Zhang et al., 2021), are estimated to have a transmission fitness effect of 1.232 (95% CI: 1.14-1.30) and 1.211 (95% CI: 1.13-1.28), respectively. B.1.1.7 has the second largest transmission fitness effect of 1.318 (95% CI: 1.27-1.35). B.1.526, which has spread rapidly in New York and carries the suspected antigenic escape mutation Spike E484K, is estimated to have the largest estimated transmission fitness effect (MLE: 1.397, 95% CI: 1.35-1.49) of all currently circulating lineages.

The fitness advantage we estimate for B.1.1.7 is much smaller than the 50-70% increase in transmissibility estimated for B.1.1.7 in the UK (Davies et al., 2021; Volz et al., 2021). However, in the US, B.1.1.7 did not have the same explosive growth as it did in the UK and remained at relatively low frequencies in early 2021 despite arriving in the US as early as October or November, 2020 (Larsen and Worobey, 2021; Washington et al., 2021). Nevertheless, to ensure our phylodynamic model is not underestimating the fitness of B.1.1.7, we also estimated the fitness of B.1.1.7 from a phylogeny of 30,000 SARS-CoV-2 genomes sampled in England between Sept 1st, 2020 and Feb. 1st, 2021. In England, we estimate the transmission fitness effect of B.1.1.7 to be 1.634 (95% CI: 1.61-1.65) relative to B.1, on par with earlier estimates (Supp. Figure 7). Thus, using the parental lineage B.1 as a basis for comparison, we estimate that the fitness of B.1.1.7 is 63% higher than B.1 in England but only 32% higher in the US.

Moreover, if anything, B.1.1.7 and other newly emerging variants are likely over-represented in the GISAID database due to preferential sequencing of variants of concern/interest. In particular, it is suspected that B.1.1.7 and other lineages with the Spike ΔH69/V70 deletion mutation are preferentially selected for sequencing in the US because this deletion leads to Spike gene target failure (SGTF) during diagnostic qPCR testing (Washington et al., 2020). Systematic oversampling of SGTF-associated variants with ΔH69/V70 could severely bias our phylodynamic fitness estimates. Indeed, a sensitivity analysis shows that the estimated fitness of B.1.1.7 declines exponentially as the assumed sampling fraction of B.1.1.7 increases (Supp. Figure 8).

Because other lineages besides B.1.1.7 share the ΔH69/V70 deletion and are also likely oversampled, we consider models that allow lineages with ΔH69/V70 to have their own SGTF-specific sampling fraction. SGTF-specific sampling fractions were estimated based on the number of GISAID sequences with the ΔH69/V70 deletion relative to the total number of COVID infections caused by a ΔH69/V70 variant. The later was imputed based on the number of SGTF-positive samples relative to all positive COVID tests using Helix’s nation-wide diagnostic qPCR testing data (see Methods). Nationally, we estimate that SGTF samples were oversampled 4.11 fold, although there is extreme spatiotemporal heterogeneity (Supp. Figure 9). Regionally, we estimate that SGTF variants were oversampled by less than 4 fold in most regions, but there were larger than 10 fold sampling biases in the Southeast (Region 4) and Southwest (Region 9).

Accounting for SGTF-specific sampling fractions in our phylodynamic model did not substantially alter the estimated fitness of PANGO lineages (Figure 6). For lineages with the the ΔH69/V70 deletion mutation, we estimate slightly lower transmission effects for B.1.427/B.1.429 but observe no change for B.1.1.7, suggesting that our fitness estimates are largely robust to SGTF-specific sampling biases.

### Decomposing the sources of fitness variation

Finally, we fit a model that included genetic features, spatiotemporal effects and branch-specific random effects to account for additional fitness variation not attributable to any feature or modeled source of variation in the model. Fitting this model to the SARS-CoV-2 phylogeny yields a fitness mapping function that we can use to predict the fitness of all lineages in terms of their transmission rate (Figure 7).

**Figure 7.**
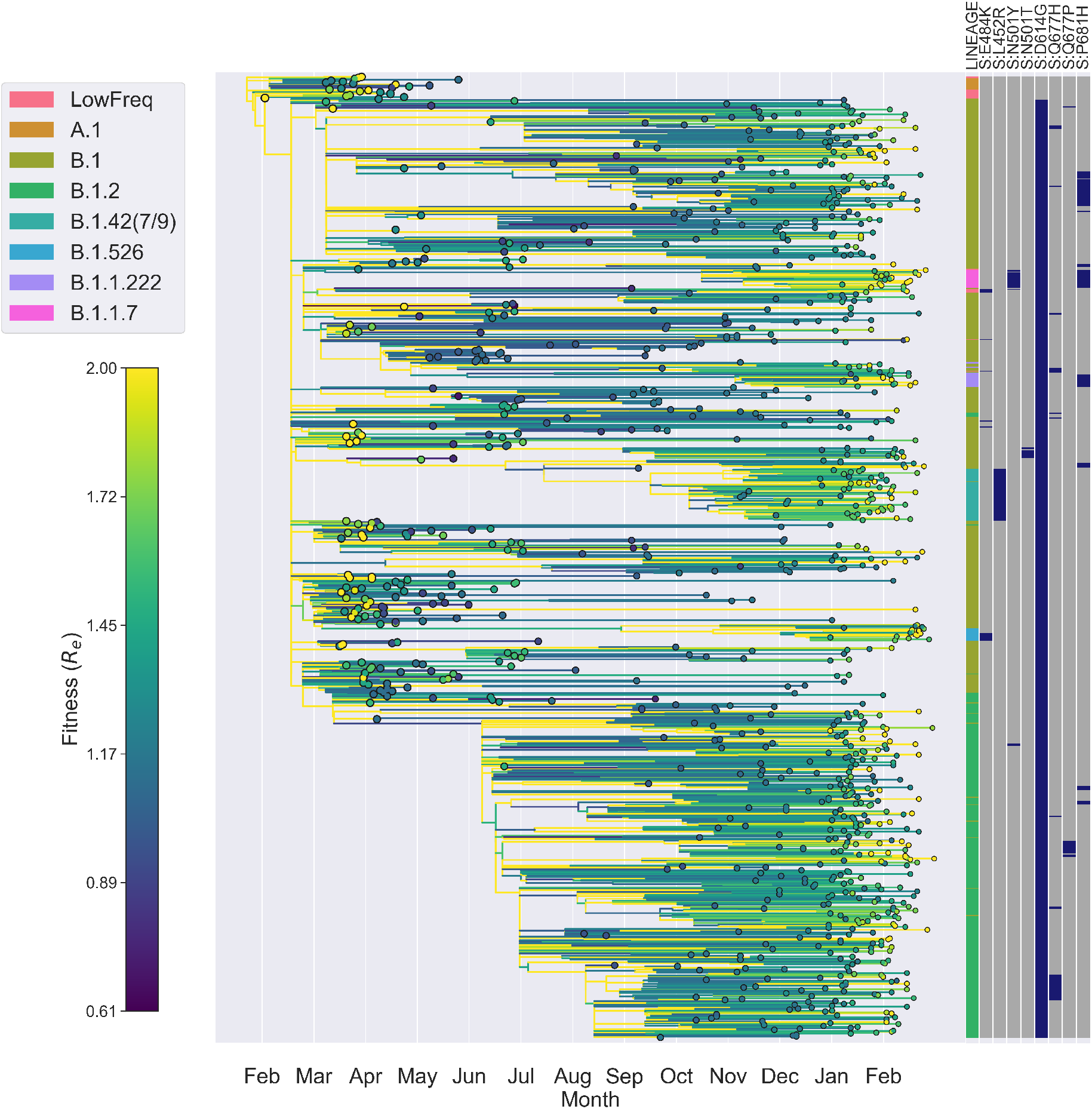
SARS-CoV-2 phylogeny with lineages colored by their transmission fitness (*R*_*e*_). The annotation block provides each sampled tip’s PANGO lineage assignment and genotype for several “landmark” genetic features in Spike. Fitness is predicted based on each lineage’s ancestral features using the fitted fitness mapping function with spatial, genetic and random effects. For the purposes of visualization, the full ML tree was thinned to include only 1000 randomly sampled tips and the fitness color scale was capped at a *R*_*e*_ = 2 to emphasize variation in fitness surrounding the mean rather than the full range of fitness values.

Given the fitness of each lineage, we can compute how much fitness varies between lineages and then decompose total fitness variation into parts attributable to different components of fitness (Figure 8). At the beginning of the pandemic, virtually all fitness variation is attributable to spatial heterogeneity in transmission among geographic regions or to random effects which cannot be explained by spatial or genetic features in our model (Figure 8B). As expected, genetic variants explain little to no fitness variation at the beginning of the pandemic when the virus population was genetically homogeneous. However, the fraction of fitness variation attributable to genetic variation quickly rises and then falls with the rise of Spike D614G in spring 2020. Genetic fitness variation then rises again in late summer and plateaus in late 2020, during which time approximately 30% of fitness variation is explained by genetic variants. Genetic variation in fitness then declines in early 2021 as less fit lineages are gradually replaced by more fit variants.

**Figure 8.**
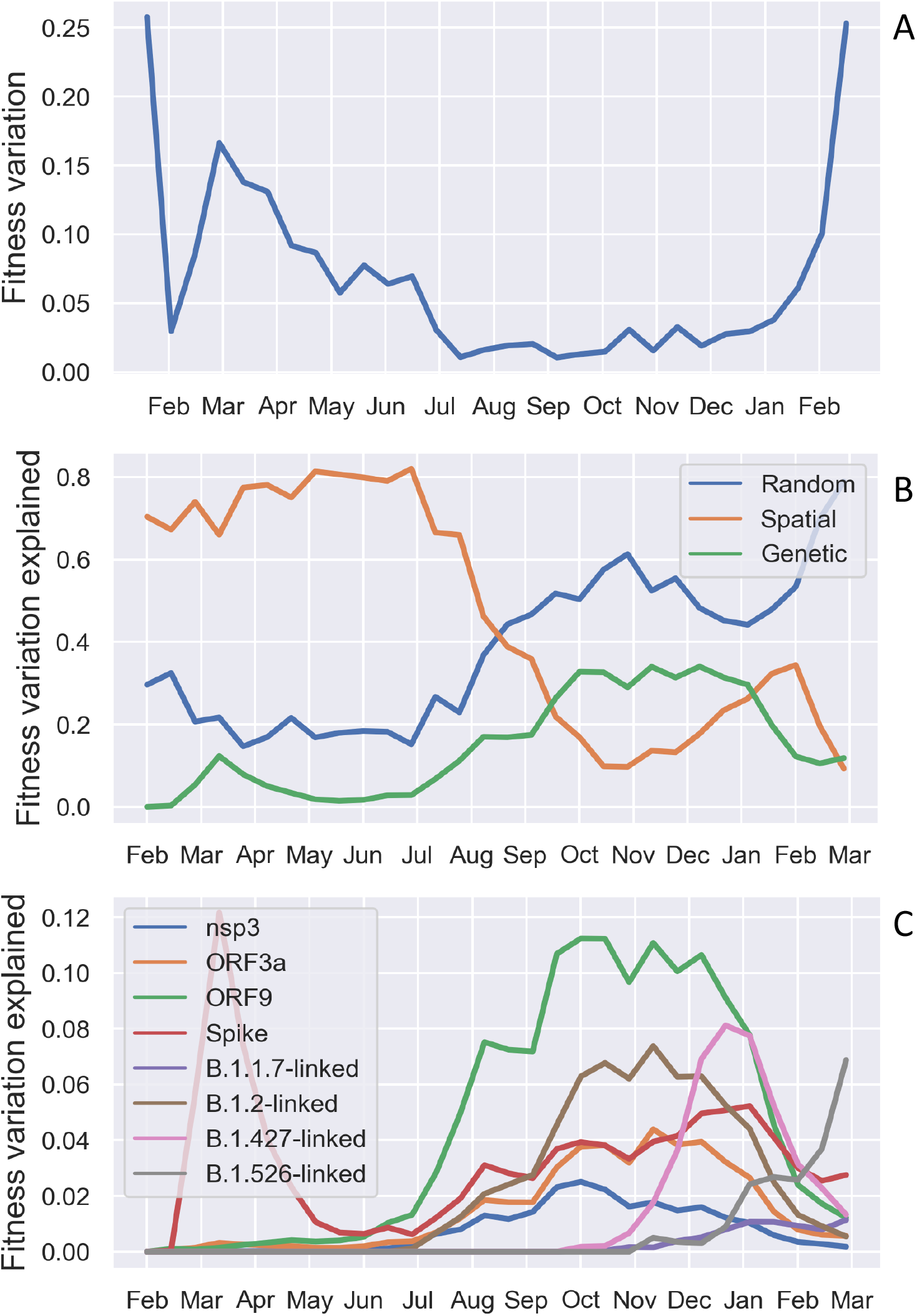
Fitness variation among lineages decomposed into sources attributable to different components of fitness. (A) Overall variation in fitness among lineages in the SARS-CoV-2 phylogeny through time. (B) Fraction of fitness variation explained by genetic, spatial and random fitness effects. (C) Fraction of fitness variation explained by different sets of genetic features.

Most genetic fitness variation is in turn attributable ta a small subset of genetic features including AAVs in Spike and sets of AAVs linked to major PANGO lineages (Figure 8C). We also included AAVs in nsp3, ORF3a, and ORF9 as we found these coding regions contain several AAVs with large positive fitness effects (Figure 3). Unexpectedly, AAVs in ORF3a and ORF9 at times contribute more fitness variation than AAVs in Spike, although this occurs mainly during the fall of 2020 when overall fitness variation is low. Nevertheless, this does suggest that less well-characterized regions of the genome outside of Spike may be shaping viral fitness in ways that remain poorly understood.

## Discussion

Determining what factors shape viral fitness variation at the between host level poses a major challenge to understanding a pathogen’s transmission dynamics more generally. We therefore developed a new phylodynamic framework for learning how a large number of genetic and non-genetic features shape the overall fitness of SARS-CoV-2 at the host population-level. A major advantage of this framework is that it allows us to decompose or partition total fitness into different fitness components to learn how both intrinsic and extrinsic factors shape viral fitness. Applying this framework to over 88,000 viral whole genomes sampled in the United States over the first year of the pandemic, our results suggest that fitness variation among lineages was largely attributable to spatial heterogeneity in background transmission rates during the first months of the pandemic. The ability to partition fitness components between intrinsic genetic and non-genetic factors even revealed that the rapid rise of Spike D614G was due, at least in part, to a large spatial transmission advantage.

However, since late 2020, an increasing fraction of fitness variation in the US is attributable to emerging variants, including the PANGO lineages B.1.1.7, B.1.526 and B.1.427/B.1.429. Although quantifying the relative fitness of these lineages is complicated by sampling biases, we estimate that these lineages have major transmission fitness advantages over earlier circulating lineages, and that our phylodynamic fitness estimates are largely robust to variant-specific sampling biases such as SGTF. In particular, we estimate that the “domestic” variants B.1.427/B.1.429 and B.1.526 have a 20 and 40% fitness advantage relative to the ancestral B.1 lineage, respectively. We also estimate that B.1.1.7 has a transmission advantage of about 32% in the US. While our estimate of B.1.1.7’s transmission advantage is not substantially lower than previous estimates of a 35-45% advantage in the US (Washington et al., 2021), these estimates are much smaller than the 50-70% transmission advantage estimated for B.1.1.7 in the UK using epidemiological data (Davies et al., 2021), coalescent-based methods (Volz et al., 2021), and our own phylodynamic birth-death methods. This smaller fitness advantage of B.1.1.7 is however consistent with its less explosive growth in the US. Despite arriving in the US as early as October or November, 2020 (Larsen and Worobey, 2021; Washington et al., 2021), B.1.1.7 remained at low frequencies in many regions until early spring 2021. B.1.1.7 may have also faced increased competition from nearly equally fit “domestic” variants, such that its growth rate in the US was impeded in an increasingly immunized population.

Another major advantage of our phylodynamic framework is that we can go beyond estimating the fitness of entire lineages and estimate the fitness effects of individual mutations while controlling for the fitness effects of other linked mutations. Exploring a large number of amino acid variants across the entire SARS-CoV-2 genome revealed moderate to large fitness effects in both expected and unexpected regions of the genome. As expected, we found several AAVs in Spike with substantial positive fitness effects on the order of 5-10%. Most of these mutations are previously described variants that either increase cellular binding avidity (D614G), escape neutralizing antibodies (L452R and E484K) or both (N501Y/T) (Deng et al., 2021; Greaney et al., 2021; Zahradnik et al., 2021). Perhaps more surprisingly, we found several AAVs in coding regions outside of Spike with large positive fitness, inclduing nsp3, ORF3a and ORF9. While these proteins remain less well-studied than Spike, they nonetheless play important roles in the viral life cycle or host-virus interactions. Nsp3 is the largest protein encoded by the SARS-CoV-2 genome and functions both as a protease and in anchoring the viral replication/transcription complex to cellular membranes (Lei et al., 2018). ORF3a is a multifunctional protein involved in cell membrane trafficking, host innate immune responses and apoptosis (Issa et al., 2020). Among bat and other non-human betacoronaviruses, ORF3a was found to have the greatest number of sites under (positive) episodic diversifying selection outside of Spike and the nucleocapsid, suggesting that it may facilitate host adaptation across species (MacLean et al., 2020a). ORF9 encodes the nucleocapsid (N), a key structural protein that is also immunogenic, although it is unclear if antibodies that recognize epitopes in N provide any neutralization potential (Gao et al., 2015; Ladner et al., 2021).

There is now considerable evidence that the Spike D614G mutation significantly alters viral fitness within individual hosts. Controlled experiments show that the D614G mutation allows the Spike glycoprotein to increase its binding affinity to the human ACE2 receptor, more than doubling cellular infectivity and viral replication rates in both cell culture and hamsters (Korber et al., 2020b; Plante et al., 2020). Higher viral replication rates could in turn explain why individuals infected with the 614G variant tend to have slightly higher viral loads (Wölfel et al., 2020; Korber et al., 2020b; Volz et al., 2020). Nevertheless, we estimate that D614G increases transmission fitness at the host-population level by only about 10%. Spike D614G therefore serves as an interesting case study to explore discrepancies in fitness estimated at the within and between scales. First, we note that increased replication rates and viral loads may not directly translate into increased infectiousness or transmission rates between hosts. While the relationship between viral load and infectiousness remains poorly understood for most respiratory viruses including SARS-CoV-2, recent work modeling clinical viral load data suggests that infectiousness does not increase linearly with viral load (Ke et al., 2020; Wölfel et al., 2020). Instead infectiousness increases with the logarithm of viral load such that it saturates at higher viral loads. This appears to fit a general pattern, as the amount of exhaled virus also saturates with increasing viral load for other seasonal coronaviruses (Leung et al., 2020b). Given that the ancestral 614D variant was already able to efficiently replicate to high viral loads (Wölfel et al., 2020), it is conceivable that any additional replication advantage provided by the 614G variant would not significantly increase transmission rates further. The enhanced cellular infectivity and replication rates of D614G within hosts is therefore not irreconcilable with our inference that the mutant had more modest population-level fitness effect.

It does initially seem more challenging to reconcile the small estimated fitness advantage of D614G with its rapid spread and near universal rise in frequency around the world, especially as our simulations show that such a moderate fitness advantage would have been insufficient to explain the explosive growth of 614G observed in the US. Our phylogenetic analysis may partially explain this discrepency, at least in the US. The ancestral 614D variant was largely limited to the West Coast (Regions 9 and 10), whereas the 614G variant established early in the Eastern US, especially in New York and New Jersey (Region 2). Due to the disproportionately large number of infections in New York and New Jersey during the early stages of the pandemic, the overall prevalence of the 614G variant also increased rapidly. This scenario is strongly supported by our phylodynamic analysis, which revealed that lineages carrying the 614G variant had a higher average background fitness due to occurring in regions with higher transmission rates as well as a genetic background with slightly higher fitness. While the 614G variant subsequently increased to high frequencies in all geographic regions, this rapid increase can partially be explained by more lineages carrying the 614G variant being imported into each region (Supp. Figure 6). A rapid increase in the frequency of the 614G variant over the 614D variant therefore does not necessarily imply that the 614G variant had a large inherent transmission advantage. In fact, when comparing the relative branching rate of lineages with either variant as a surrogate for transmission rates, we find that 614G only had about a 13% higher transmission advantage than 614D, supporting our phylodynamic estimates of an approximatre 10% fitness increase.

Borrowing the concept of gene surfing from spatial population genetics may help to explain the rapid rise of the D614G variant. Gene surfing describes a scenario where a mutation can rapidly expand its geographic range by occurring along the edge or wave front of a spatially expanding population and then “surf” to high frequencies by riding the wave of spatial expansion (Edmonds et al., 2004; Klopfstein et al., 2006; Hallatschek and Nelson, 2008). While perhaps not a perfect metaphor here because SARS-CoV-2 did not spread as a spatially cohesive wave across the US, the gene surfing analogy captures the idea of how even a neutral mutation can be propelled to high frequencies across a range of spatial locations as a result of rapid population expansion. Viewed from this prospective, one can see why spatially aggregated time series of variant frequencies can be positively misleading about the fitness of a variant during a rapidly spatially expanding epidemic. Phylogenetic analysis coupled with ancestral state reconstructions offer a means of avoiding these pitfalls because they allow us to first identify lineages in the same transmission environment (e.g. geographic region) and then quantify the relative transmission rate of lineages from their branching pattern in the phylogeny.

While our phylodynamic inference framework accounts for many potentially confounding factors including background variation in transmission rates, our analysis still has a number of limitations. First, inferences of pathogen fitness from phylogenies will inevitably depend on what lineages are sampled and included in the phylogeny. We can partially correct for sampling biases using time, location and even variant-specific sampling fractions but our fitness estimates will be dependent on the assumed sampling fractions. Second, we chose a simple fitness mapping function that assumes each feature has a multiplicative effect on lineage fitness (such that log fitness is an additive linear function of features). In reality, the relationship between a pathogen’s genotype, environment and other features may be considerably more complex due to nonlinear relationships between features and fitness or interactions among genetic features (epistasis) and the environment (GxE interactions). It is therefore likely that some of the fitness variation attributed to random effects under our model are actually due to additional genetic sources such as epistatic interactions among mutations that cannot be captured under our simple multiplicative fitness model. Learning what types of functions are expressive enough to capture these complexities while remaining statistically tractable and biologically interpretable is a major challenge for future work. Finally, the computational efficiency of our approach relies on first reconstructing phylogenies and ancestral states before fitting our phylodynamic birth-death model. While we partially account for phylogenetic uncertainty by fitting models to replicate bootstrap phylogenies, using pseudoreplication to account for uncertainty is certainly a large step back from fully Bayesian phylodynamic methods that jointly infer key evolutionary and epidemiological parameters while simultaneously integrating over phylogenetic histories. New inference methods are clearly needed to fit complex phylodynamic models to genomic data sets as large as those currently available for SARS-CoV-2.

As natural and vaccine-induced immunity continues to rise in the population, new antigenic escape mutations and other variants will likely continue to arise. Our phylodynamic framework can used to examine the epidemiological significance of such mutations by estimating their transmission potential while accounting for confounding sources of fitness variation. Another major advantage of our approach is that it allows us to learn the relative importance of different features to overall pathogen fitness by decomposing fitness variation into its component parts. In the future, this will allow us to determine the contribution of new genetic variants relative to extrinsic factors such as host mobility or immunity. Because fitness variation at the host population-level is essentially equivalent to variation in transmission potential, learning what features contribute the most to fitness variation is tantamount to learning what features most strongly regulate transmission. Thus, our phylodynamic learning framework not only allows us to estimate fitness, but understand what components of fitness shape both the evolutionary and epidemiological dynamics of viral pathogens.

## Models and Methods

### General approach

Our primary goal is to learn how multiple different character traits or *features*, which may include genetic variants, phenotypic traits and environmental variables, all act together to determine the fitness of pathogen lineages in a phylogenetic tree. We assume here that the phylogeny as well as ancestral features corresponding to the ancestral state of each feature is reconstructed beforehand. The relationship between predictive features and fitness is modeled using a *fitness mapping function* that predicts the expected fitness of a lineage based on its reconstructed ancestral features. The fitness mapping function can then be used to compute the expected fitness a lineage in terms of its birth and/or death rate. For a pathogen phylogeny, birth events are assumed to correspond to transmission events and deaths correspond to recovery or removal from the infected population. Given the birth and death rates of each lineage in a phylogenetic tree, the likelihood of the tree evolving as observed can be computed analytically under a birth-death-sampling model (Stadler, 2009; Barido-Sottani et al., 2018). Our problem therefore reduces to finding the fitness mapping function that maximizes the likelihood of the phylogeny given the ancestral features of all lineages in the tree.

### Phylogenetic reconstruction

For the original Pre-2020-09 data set, a total of 22,416 SARS-CoV-2 whole genome sequences from the United States were downloaded from GISAID (Elbe and Buckland-Merrett, 2017) on October 2nd, 2020 representing sequences that were sequenced prior to September 1st, 2020. Genomes were aligned against a reference genome (NC 045512.2) using MAFFT version 7.475 (Katoh and Standley, 2013). A maximum likelihood (ML) phylogenetic tree was reconstructed in RAxML (Stamatakis, 2014) using the rapid bootstrapping method with 10 bootstrap replicates assuming a GTR model of sequence evolution with Gamma-distributed rate variation among sites. The best ML and all bootstrapped trees were then dated using LSD (To et al., 2015) assuming a fixed clock rate of 0.0008 substitutions per site per year. A total of 93 sequences were discarded due to inconsistencies in sampling times or poor sequence quality. Due to the large number of viral samples in the Post-2020-09 data set, we extracted a maximum likelihood (ML) phylogeny from a precomputed global SARS-CoV-2 ML phylogeny provided by Rob Lanfear’s group on GISAID (Lanfear, 2020). For all analyses conducted here, we use the 2021-03-13 version of the global tree. A focal tree containing all 66,339 viruses sampled in the US between September 1st, 2020 and March 1st, 2021 was then extracted using the *extract tree with taxa* function in Dendropy version 4.5.1 (Sukumaran and Holder, 2010). The extracted ML tree was then dated using LSD (To et al., 2015). For the fitness analysis of B.1.1.7 in the UK, we randomly sampled 30,000 viral isolates sampled in England between September 1st and February 1st. A focal tree containing these samples was then extracted from the global global SARS-CoV-2 ML phylogeny.

### Ancestral state reconstruction

In the original analysis of the Pre-2020-09 data, ancestral states were reconstructed for each feature under a continuous-time Markov chain model of trait evolution using PastML (Ishikawa et al., 2019). PastML estimates the relative transition rate between each pair of states and the global (absolute) rate at which transitions occur. The relative transition rates are constrained to be proportional to the equilibrium frequencies of each state as under a F81 model of nucleotide substitution. Rate parameters were estimated independently for each feature. At each internal node, the state with the highest marginal posterior probability was taken to be the ancestral state for a given feature.

For the larger Post-2020-09 data set, we reconstructed ancestral states using maximum parsimony (MP) to expedite analysis. This was motivated by the observation that there was typically little uncertainty surrounding the ML ancestral state reconstructions performed by PastML. For most genetic features, there was only a single or a small number of state transitions across the entire tree. In this case, MP and ML reconstructions are expected to agree as the most parsimonious and most likely reconstructions will be consistent given a slow rate of character evolution (Tuffley and Steel, 1997; Zhang and Nei, 1997). MP reconstructions were performed using Sankoff’s dynamic programming algorithm (Sankoff, 1975) implemented in Python (see AncestralReconstruction.py).

Reconstructed ancestral features were then combined into a vector of categorical variables *x*_*n*_ for each lineage *n*. For categorical variables with more than one state, we used one-hot binary encoding to yield a strictly binary feature vector. Ancestral features were reconstructed for each bootstrap phylogeny independently.

### Fitness mapping functions

Our main goal is to learn the fitness mapping function *F*(*x*_*n*_) that maps the features of a lineage *x*_*n*_ to that lineage’s expected fitness. While *F*(*x*_*n*_) could be any arbitrary function, we use a simple model that assumes the fitness effect *β*_*i*_ of each feature *i* is multiplicative:

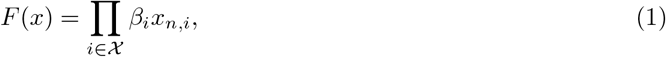

where 𝒳 is the set of all features used to predict fitness. Each feature *x*_*n,i*_ is assumed to be encoded as a binary variable or as the probability of the lineage having a particular feature.

In order to decompose fitness into its component parts below we consider fitness effects on a log scale, which gives us the additive linear model:

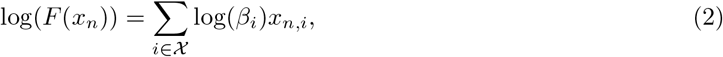

We also consider a fitness mapping function with random, branch-specific fitness effects *u*_*n*_:

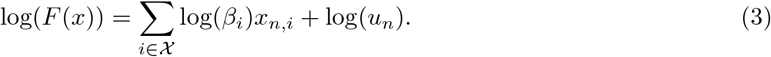

These random effects capture unmodeled sources of fitness variation such as genetic background effects at loci not included as features in the model.

Estimating branch-specific random fitness effects without additional constraints leads to extreme variability in fitness among lineages. In particular, long branches are estimated to have low fitness and short branches are estimated to have high fitness as this maximizes the likelihood of each branch under a birth-death model. We therefore use a Brownian motion model of trait evolution that constrains the branch-specific random fitness effects to be correlated between parent and child branches. Because each branch is assumed to have a unique random effect, we only allow fitness to change at birth/transmission events in the tree. The probability of a child having random fitness effect *u*_*c*_ given its parent’s random fitness effect *u*_*p*_ is

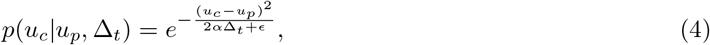

where Δ_*t*_ is the time elapsed between the parent and child node and *α* scales the variance of the child’s fitness distribution. For numerical stability, we include a small value *ε* to ensure the probability does not become infinitely small when *σ* Δ_*t*_ *<<* 1.0. This model is conceptually similar to the ClaDS model of Maliet et al. (2019) which estimates lineage-specific diversification rates by allowing for small shifts in birth and/www.w3.org/TR/xht branching events, although the CLaDS model assumes a log-normal fitness distribution for child lineages independent of branch lengths.

How much fitness is allowed to vary between parent-child lineage pairs due to random effects is controlled by the hyperparameter *α*. We estimate *α* using k-fold cross-validation. Inspired by cross-validation techniques for time series data (Roberts et al., 2017), we longitudinally cross-section or block phylogenetic trees into training and test intervals. Random fitness effects are estimated for each branch in the tree during the training interval and then lineages in the test interval inherit their random fitness effects from their parent (or most recent ancestor) in the training interval. Thus, if the random fitness effects capture true fitness variation among lineages in the training interval, these fitness effects should more accurately predict the fitness of descendent lineages and improve the likelihood of the phylogeny in the test period. In contrast, a model with *α* set too high will overfit the fitness variation among lineages in the training period but will not improve performance in the test period. We can therefore use cross-validation to estimate an optimal value of *α* that maximizes the likelihood of trees in the test period while preventing the random fitness effects from overfitting fitness variation among lineages.

### The phylodynamic birth-death-sampling model

The likelihood of a phylogenetic tree evolving as observed can be computed under a phylodynamic birth-death-sampling model (Stadler, 2009) given the expected fitness of each lineage, which we predict based on a lineage’s ancestral features *x*_*n*_ using a fitness mapping function *F* (*x*_*n*_). We assume throughout that fitness is directly proportional to a lineage’s birth or transmission rate *λ*_*n*_ = *f*_*n*_*λ*_0_, where *λ*_0_ is a base transmission rate which is scaled by a lineage’s fitness *f*_*n*_. We also assume that the removal rate *µ* and sampling fraction *σ* are constant across all lineages, although we consider models where *σ* is allowed to vary by time and region below. This dramatically simplifies the model, as instead of having a multi-type birth-death process we have a series of connected single-type birth-death processes along lineages who’s birth and death rates are piecewise constant.

Under this model, it is possible to analytically compute the likelihood of the phylogeny evolving as observed, allowing for efficient statistical inference. Given the birth, death and sampling rates and the fitness mapping function to compute the expected fitness of each lineage, the likelihood of each lineage or subtree evolving is independent conditional upon knowing the ancestral features used to predict fitness. The total likelihood of a phylogenetic tree 𝒯 can be decomposed into the likelihood of a set of sampling events *S*, a set of branching (transmission) events *B*, and a set of lineages *N* :

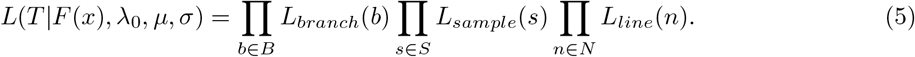

The likelihood of an individual branching or transmission event is:

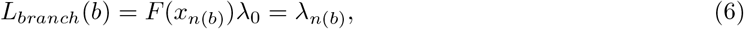

where we use the notation *n*(*b*) to refer the parent lineage involved in a paritcular branching event *b*.

The likelihood of an individual sampling event at time *t* in the past is:

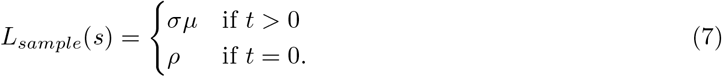

Before the present, the probability of a sampling event depends on the removal rate *µ* and the probability *σ* that the lineage is sampled upon removal. At the present (*t* = 0), any extent (i.e. currently infected) individual is sampled with probability *ρ*.

*L*_*line*_(*n*) gives the likelihood a lineage *n* evolved as observed; i.e. the probability that the lineage survived without giving rise to other sampled lineages. As shown in Barido-Sottani et al. (2018), over a time interval of length  Δ_*t*_, this likelihood can be computed as:

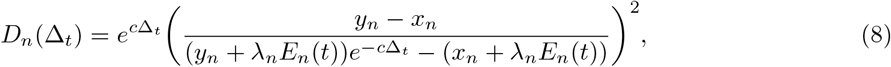

with:

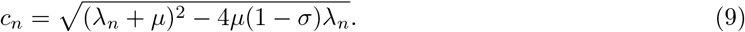

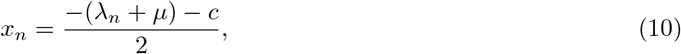

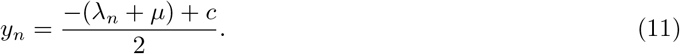

The *E*_*n*_(*t*) terms in (8) represent the probability that a lineage at time *t* in the past produced no sampled descendants. Assuming that the birth, death and sampling rates do not change along unsampled lineages from their values at time *t*, these probabilities are given by:

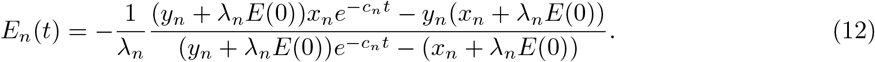

*E*(0) is the initial condition or probability of lineage not being sampled at the present (*t* = 0). Given that the proportion of individuals sampled at present is *ρ*, we set *E*(0) = 1 − *ρ*. For simplicity we assume that *ρ* = *σµ/*365 so that the probability of a lineage being sampled on the final day of sampling is proportional to the probability of an individual being removed from the infectious population on that day, but the sampling fraction is the same as any point in the past.

### Model fitting and statistical inference

Learning the fitness mapping function from a phylogenetic tree is a somewhat non-standard problem in that we do not have direct observations of a lineage’s fitness to which we can compare our predictions under *F* (*x*). Nevertheless, we can formulate statistical inference as an optimization problem where we seek to find the fitness mapping function *F* (*x*) with parameters 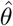 that maximizes the overall likelihood of the phylogeny given the reconstructed ancestral features under the birth-death-sampling model:

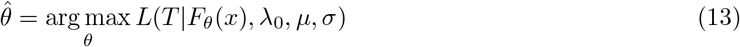

Formulating the problem in this way opens the way to using efficient optimization algorithms developed in recent years to train neural networks and other machine learning models. Instead of optimizing a typical loss function (e.g. least-squares), we simply maximize the likelihood of the phylogeny under the birth-death-sampling model. In particular, we use the ADAM optimizer (Kingma and Ba, 2014), a form of stochastic gradient descent (SGD) which adapts learning rates based on gradients (i.e. first-order derivatives) of the likelihood function with respect to different parameters. Adapting the learning rates allows the algorithm to accelerate its momentum towards parameters that optimize the loss function. To make use of ADAM and other high-performance SGD algorithms, we implemented our fitness mapping function and birth-death likelihood function in TensorFlow 2 (Abadi et al., 2016). Gradients in the likelihood function are computed using TensorFlow’s auto-differentiation functionality, allowing us to efficiently fit complex models with hundreds of features or parameters. Using this approach, even fitting our most complex model with over 300 free parameters to a phylogeny with over 22,000 tips only takes a few minutes on a standard desktop computer.

Learning the fitness mapping function through gradient descent provides maximum likelihood estimates (MLEs) of each parameter in the model. To quantify uncertainty surrounding the MLEs, we compute the likelihood of the phylogeny over a fixed grid of parameters values and then determine which values fall within the 95% credible intervals using an asymptotic chi-square approximation to the likelihood ratio test.

### Performance on simulated data

To test the ability of our methods to correctly estimate fitness effects, we ran forward simulations where both genetic and spatiotemporal features influence viral fitness. Phylogenies were simulated under a birth-death-sampling model using the stochastic Gillespie algorithm (Gillespie, 2007) starting with a single infected individual. In all simulations we assume a constant base birth rate of 1.2 and death rate of 1.0 per time unit. A virus’s genotype is represented by ten binary sites where zeros indicate the ancestral state and ones indicate the mutant state. Each site has a random, multiplicative effect on fitness when mutated to the one state. Mutation occurs at a constant per site rate of 1.5 x 10^*−*2^ per time unit. A lineage’s spatial location is encoded as an additional evolving character trait. To emulate background fitness variation due to spatiotemporal heterogeneity in transmission, each combination of region and time interval is assigned a background transmission rate. Individuals move from one region to another with a transition rate of 0.3 per time unit. Furthermore, in order to emulate an additional source of fitness variation not directly accounted for in the inference model, we added transmission heterogeneity by having each infected individual draw a random effect that rescales their transmission rate from a gamma distribution (Lloyd-Smith et al., 2005). Here the gamma distribution has a dispersion parameter 0.15 and scale parameter 10, such that on average, there is a branch-specific fitness of 1.5, but individually, fitness varied substantially.

Performance was tested under both a high sampling regime (*σ* = *ρ* = 0.5) and a low sampling regime (*σ* = *ρ* = 0.05). A phylogeny was built from the true ancestral history of sampled individuals. True ancestral features (states) were assumed to be known for the purposes of validating the inference algorithm. Simulations were run for 8 time units and those that ended more than 0.2 time units before then, or that had less than 800 sampled individuals, were discarded. We then estimated background transmission rates and genetic fitness effects from each simulated phylogeny.

Estimated spatiotemporal and genetic fitness effects are generally well correlated with the true values used in simulations (Supp. Figure 10). However, estimation accuracy depends largely on the overall sampling fraction and the number of individuals sampled with a given feature (spatial location or genotype). In particular, the fitness effects of rare features sampled at low frequencies tend to have the most variable and least accurate estimates. Because estimating the fitness of rare features under a birth-death model appears to be inherently difficult (Rasmussen and Stadler, 2019), we only estimate fitness effects for features with a sampling frequency above 0.5% from empirical SARS-CoV-2 phylogenies.

### Birth-death-sampling model parameters for SARS-CoV-2

Because it is not possible to estimate all of the parameters in the birth-death-sampling model from a phylogeny alone, we fix some parameters at values based on prior knowledge. We assume individuals infected with SARS-CoV-2 stay infected (and infectious) for 7 days on average, leading to a removal rate 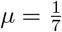 per day.

In several models we allow the base transmission rate *λ*_0_ or sampling fraction *σ* to vary over time. In this case we have a time-varying transmission rate *λ*(*t*) and *σ*(*t*) that depends on the time *t*. However, this can easily be incorporated into the birth-death model above. If a lineage’s transmission rate or sampling fraction changes along a branch due to an underlying change in *λ*(*t*) or *σ*(*t*), we simply divide the branch into segments corresponding to the time intervals over which these parameters remain piece-wise constant and add each lineage segment to the set of lineages in *N*.

We assume that the sampling fraction *σ* was zero before the first sample in our data set was collected in January 2020. After the first sampling date, we allow the sampling fraction to vary by time and region as described below.

### Modeling sampling heterogeneity

In order to estimate how sampling fractions vary across space and time, we count the number of sequence samples *g*_*i,t*_ submitted to GISAID within each geographic location *i* over each time interval *t*. An unbiased estimate of the sampling fraction *σ*_*i,t*_ would therefore be:

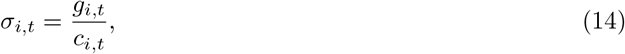

where *c*_*i,t*_ is the total number of infections or cumulative incidence in region *i* over time interval *t*.

We of course do not know *c*_*i,t*_ but can obtain a pseudo-empirical estimate *ĉ*_*i,t*_ by considering the number of deaths attributed to SARS-CoV-2 *d*_*i,t*_ and the estimated infection fatality ratio *φ*, which was assumed to be 0.5% (Perez-Saez et al., 2021). We can therefore approximate the total number of cases *c*_*i,t*_ as:

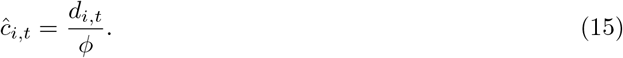

Substituting *ĉ*_*i,t*_ for *c*_*i,t*_ in (14), we arrive at a crude estimate of the sampling fraction.

While the case fatality ratio likely also fluctuates over space and time due to changes in the age distribution of infections among other reasons, it seems reasonable to assume that the mortality rate fluctuates less than the testing or sequence sampling fraction (Flaxman et al., 2020). We can therefore roughly estimate the total number of cases based on the number of observed deaths. Using this approach, we estimate that there were a total of 35,134,400 cumulative cases in the US by September 1st, whereas the total number of positive cases reported by the COVID Project (https://covidtracking.com/data/national) on the same date was 6,017,826. Our estimate for the total number of cases suggests that 83% of all infections were not detected in the US, which is consistent with recent estimates by Wu et al. (2020), who estimated that up to 89% of all infections are unreported using an independent approach.

Using data from the COVID Project to tabulate cumulative deaths *d*_*i,t*_ for each region and time interval, we estimate how the sampling fraction *σ*_*i,t*_ varied across regions and time (Figure 1A). For these estimates we assume reported deaths lag behind reported cases by three weeks when estimating sampling fractions.

### Modeling variant-specific sampling biases

An additional complication arises here because B.1.1.7 and other lineages carrying the Spike  ΔH69/V70 deletion mutation are likely oversampled due to preferential sequencing of viral isolates suspected of being newly emerging variants based on Spike gene target failure (SGTF) during diagnostic qPCR testing (Washington et al., 2020). To account for SGTF-related sampling bias, we estimate a SGTF-specific sampling fraction for lineages with the ΔH69/V70 deletion. Note that we estimate sampling fractions for all lineages with the ΔH69/V70 deletion rather than just B.1.1.7 as other lineages share this deletion and are therefore likely also preferentially selected for sequencing. We estimate SGTF-specific sampling fractions based on the fraction of SGTF-positive samples 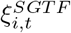 relative to the total number of SARS-CoV-2 positive samples using Helix’s nation-wide diagnostic qPCR data (https://github.com/myhelix/helix-covid19db/blob/master/counts_by_state.csv). We then modify (14) by multiplying the total number of infections *c*_*i,t*_ by the SGTF-positive fraction *ξ*_*i,t*_:

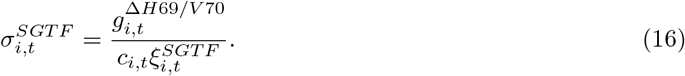

Here, 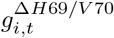 is the number of sequence samples deposited to GISAID with the Δ*H*69*/V* 70 deletion.

### Model selection

We initially fit several different models that allowed background transmission rates to vary over space and time in different ways and compared their relative fit using AIC. This model selection step was performed only on the the Pre-2020-09 data set. Compared to our base model which assumes a constant transmission rate across both space and time, a model that allows transmission rates to vary by geographic region increases the likelihood of the phylogeny and model fit as quantified by AIC (Table 4). A similar model that allowed transmission rates to vary by US state instead of region further improves model fit.

**Table 4.** Model selection using the maximum log likelihood 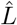 for each model and AIC

| Model | # params | $\hat{L}$ | AIC | ΔAIC |
| --- | --- | --- | --- | --- |
| Base | 1 | 15630.6 | -31259.2 |  |
| Spatial effects (by region) | 10 | 15667.6 | -31315.2 | -56.0 |
| Spatial effects (by state) | 52 | 15717.2 | -31330.4 | -71.2 |
| Temporal effects | 9 | 16655.9 | -33293.8 | -2034.6 |
| Spatial (by region) x temporal effects | 90 | 17290.1 | -34400.2 | -3141.0 |
| Spatial (by state) x temporal effects | 468 | 18083.6 | -35231.2 | -3972.0 |

Allowing transmission rates to vary over time in a piece-wise constant manner using monthly time intervals improves model fit more than allowing transmission rates to vary by location. Using biweekly rather than monthly time intervals does not improve model fit further. In turn, all models with only spatial or temporal effects are vastly outperformed by a model that allows transmission rates to vary by both time interval and geographic location (spatiotemporal effects). Using states instead of geographic regions increases the likelihood of the spatiotemporal effects model and has lowest overall AIC value, but we continue to use the model with regional spatial resolution as several states are very poorly represented in the GISAID database. We therefore allow transmission rates to vary by geographic region over monthly time intervals in all subsequent analyses.

### Decomposing fitness variation

Given the ancestral features *x*_*n*_ of a lineage, we can compute the lineage’s fitness using the fitness mapping function. We can then partition or decompose total variation in fitness between lineages into sources attributable to different components of fitness. To do this, we first partition the features in *𝒳* into different disjoint, non-overlapping subsets *𝒳*_*k*_ *⊂ 𝒳*; *𝒳*_*k*_ *∩ 𝒳*_*l*_ = *∅* for all subsets *k* and *l*.

In the fitness mapping functions presented above, each feature *i* has a fitness effect *f*_*n,i*_ on a lineage’s fitness, where *f*_*n,i*_ = *β*_*i*_*x*_*n,i*_. We let the vector ***f***_***i***_ hold the fitness effect of feature *i* for all lineages in the phylogeny and ***f*** hold the overall fitness of each lineage in the phylogeny. Under the additive model that considers fitness on the log scale (2), ***f*** = ∑_*i*_ ***f***_***i***_. Using the general property that the variance in the sum of random variables is equal to the sum of their individual variances and covariances, we can partition the total variation in fitness into variances attributable to individual features and covariances attributable to pairs of features:

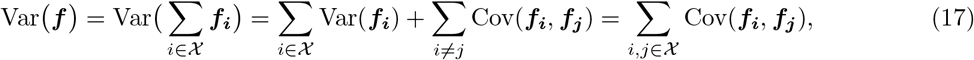

where the covariances account for the fact that the features may be correlated across lineages and therefore not independent.

We can take advantage of the additive property of the variances to compute the fraction of total variance attributable to any particular subset of features *𝒳*_*k*_:

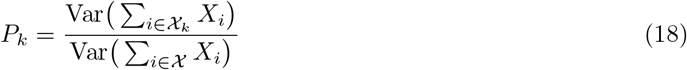

In our SARS-CoV-2 analysis, we partition features into three different components of fitness: genetic, spatial and random (unexplained) effects. To ensure that the fraction of variance attributable to each component sum to one, we compute the fraction of variation attributable to each fitness component as:

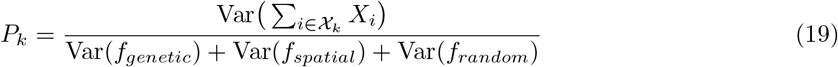

In other words, we ignore the covariances among fitness components. We do this to ensure that negative covariances among components do not cause the variance attributable to a particular component to be greater than the total variance.

### Two-strain epidemiological model

In order to explore if the transmission fitness effect of approximately 10% that we estimate for Spike D614G would have been sufficient to explain its rapid rise in frequency over the spring of 2020, we simulated the evolutionary dynamics of a mutant variant in a single host population under a two-strain Susceptible-Infected-Exposed-Recovered (SEIR) model parameterized for Covid-19. In this model, an initial resident strain (614G) with transmission rate *β* seeds the epidemic and then a mutant strain (614D) with transmission rate *β*_*m*_ = *βf*_*m*_ enters the population through external introductions. We then systematically vary the mutant’s fitness *f*_*m*_ to see how much more fit the mutant needs to be in order to match the evolutionary trajectory of D614G.

The epidemic dynamics in the host population are described by the following system of differential equations:

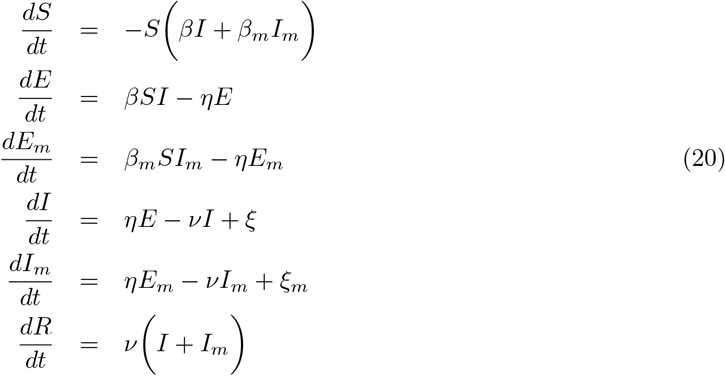

Here, *η* is the incubation rate at which exposed individuals become infectious and *ν* is the removal or recovery rate. We assume a 4 day incubation period (*η* = 0.25) and a 7 day infectious period (*ν* = 0.143) (Davies et al., 2020; Ferretti et al., 2020). *ξ* and *ξ*_*m*_ give the rate of external introductions into the host population (per capita) of the resident and mutant strain, respectively.

We assume that there was a single infected individual in the population on Jan. 15th 2020, reflecting the timing of the earliest probable infections in the US (Worobey et al., 2020). External introductions of the resident strain occur at a rate of 1 per day after Jan. 15th. The external introduction rate of the mutant is initially zero, but switches to *ξ*_*m*_ *>* 0 after Feb. 15th to reflect the earliest probable introductions of D614G into the US. Because the relative rate at which the 614D and 614G entered the US through external introductions is a key unknown that largely determines the evolutionary trajectory of the mutant, we explore different ratios of *ξ* and *ξ*_*m*_.

Finally, since no constant transmission rate can recapitulate the epidemic dynamics of Covid-19 in the US under the SEIR model, we allow the base transmission rate *β* to decline over time to mimic the effects of social distancing or other interventions. We let *β* decrease between piece-wise constant intervals such that *R*_*e*_ is 2.5 between Jan. 15th and Feb. 15th, 1.5 between Feb. 15th and Mar. 15th, 1.25 between Mar. 15th and Apr 15th and 1.1 after Apr. 15th, reflecting the average *R*_*e*_ values inferred for these time intervals under our phylodynamic model.

## Code and data availability

Code and data to replicate our phylodynamic analysis is freely available on GitHub at github.com/ davidrasm/phyloTF2.

## Acknowledgments

We would like to thank GISAID and all of the researchers who have submitted SARS-CoV-2 sequences to the GISAID database. A full list of authors and originating laboratories for GISAID submissions we use here is available at: https://github.com/davidrasm/phyloTF2/blob/main/gisaid_hcov-19_acknowledgement_table_2020_12_11_10.pdf. DAR is supported by the US Dept. of Agriculture Hatch project 1016556. LK is supported by the US Centers for Disease Control and Prevention (U01CK000587-01).

## Supplementary Figures

**Supp. Figure 1.**
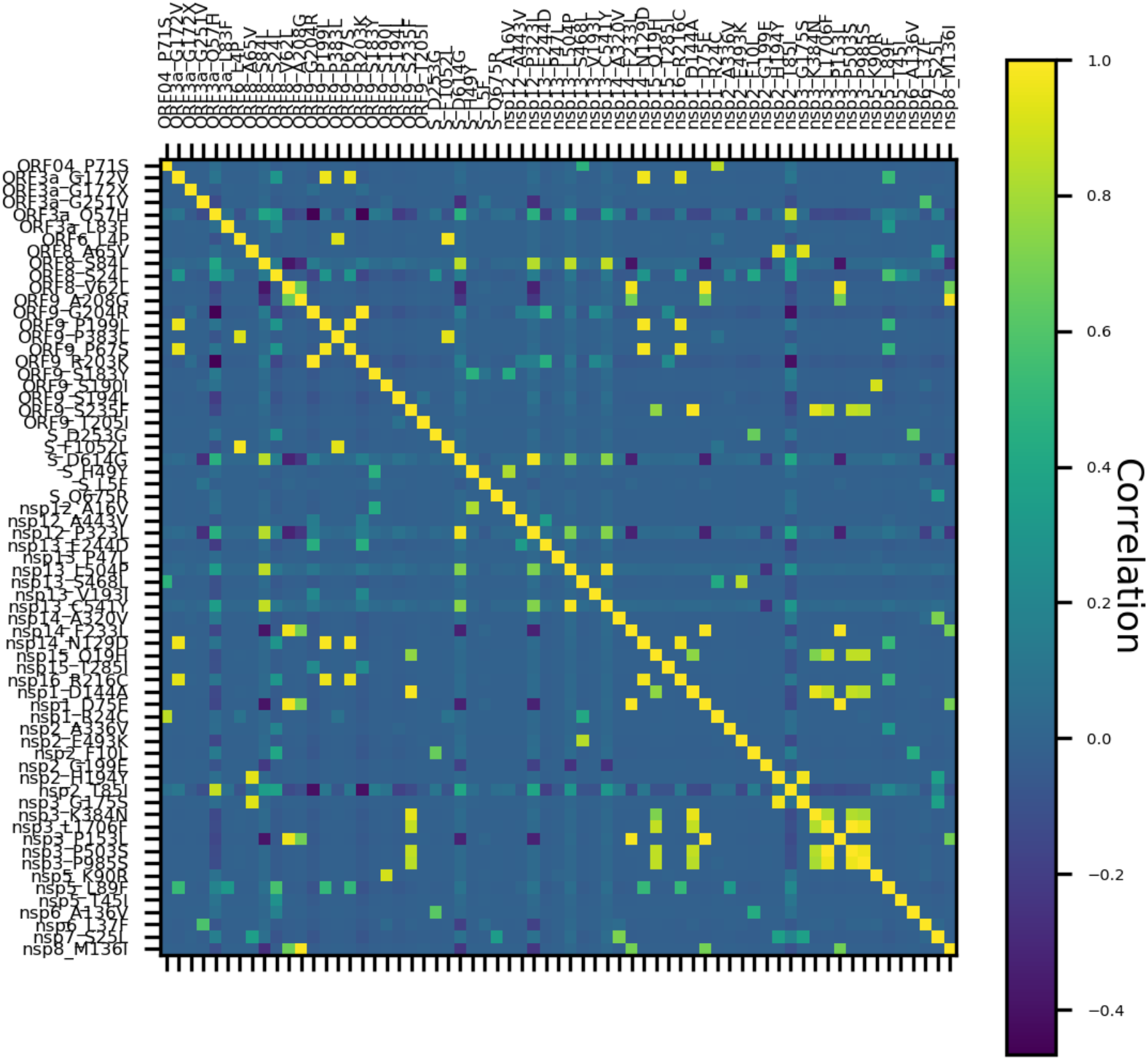
Correlation among amino acid variants in sampled SARS-CoV-2 genotypes. Colors in the heat map represent the Pearson correlation coefficient between each pair of variants.

**Supp. Figure 2.**
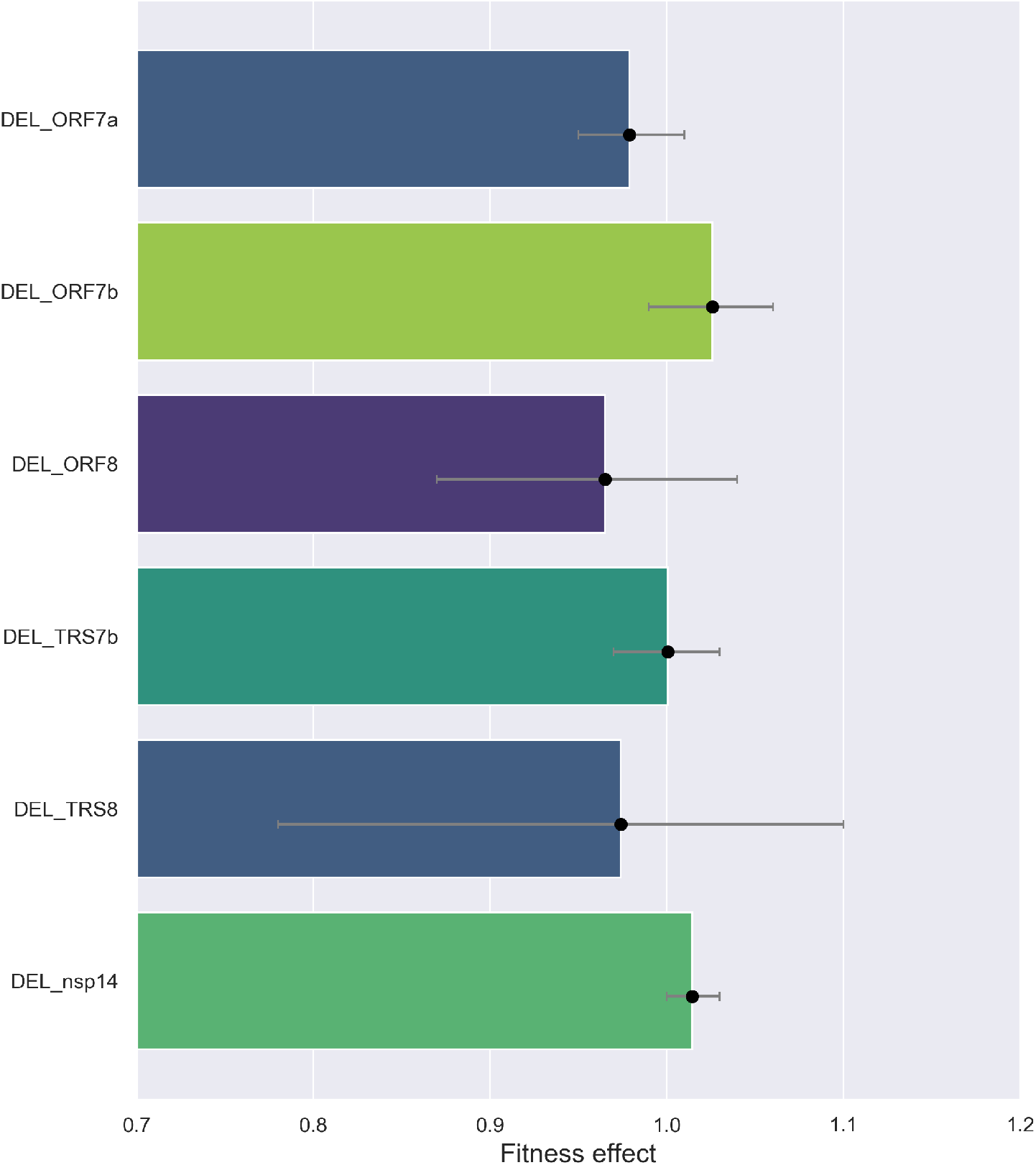
Estimated fitness effects of structural variants with major deletions. Bars are colored according to the maximum likelihood estimate of the fitness effect of each variant. Capped lines indicate 95% credible intervals. Deletion mutations in these regions tend to be hypervariable in both number and their starting and end positions. We therefore grouped all structural variants with major deletions in one of these six regions. In the reference Wuhan-Hu-1 SARS-CoV-2 genome, these deletions correspond to positions: 27397-27736 (DEL ORF7a), 27759-27891 (DEL ORF7b), 27897-28262 (DEL ORF8), 27708-27752 (DEL TRS7b), 27884-27897(DEL TRS8) and 19276-19579 (DEL nsp14).

**Supp. Figure 3.**
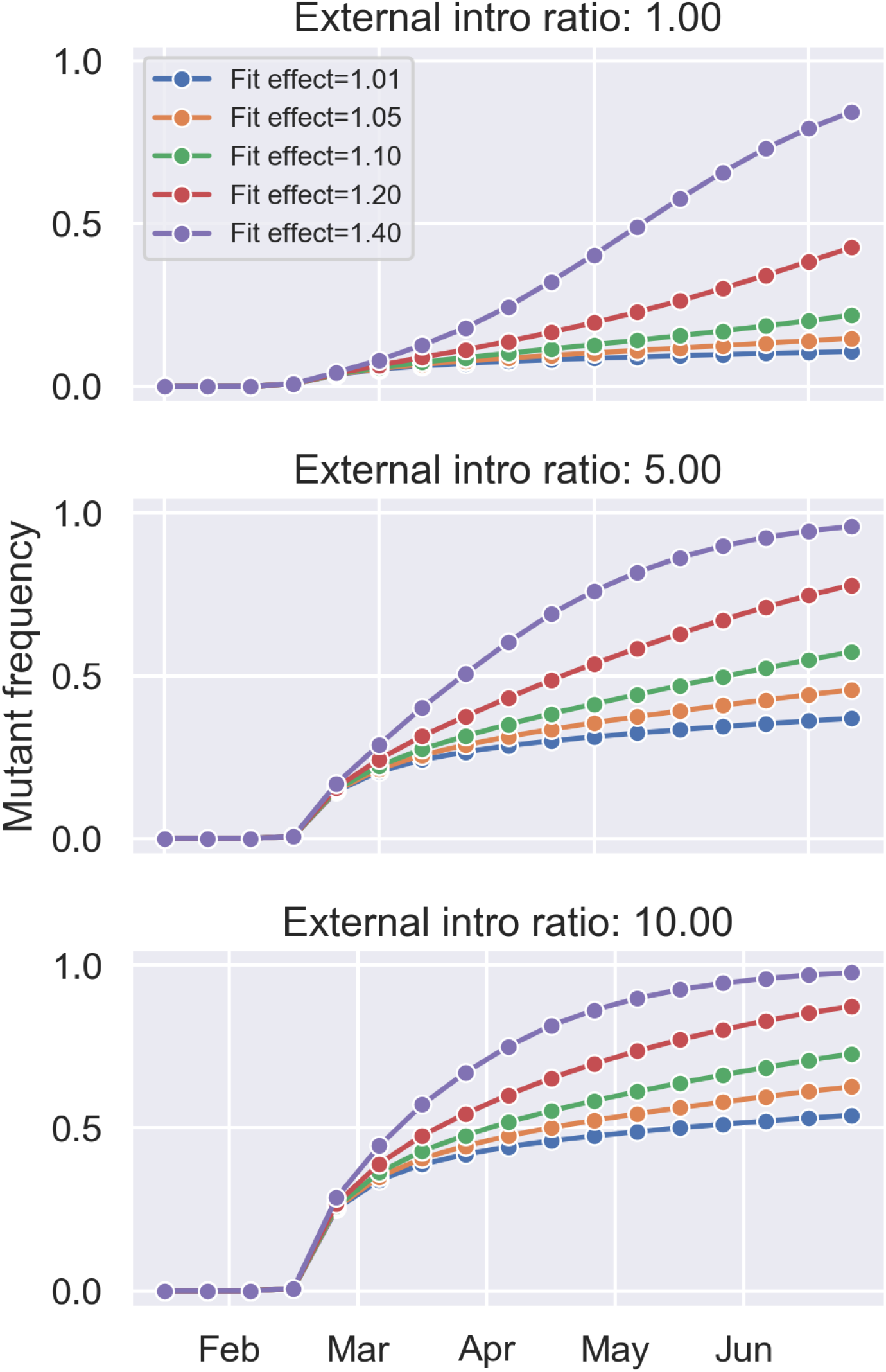
Simulated evolutionary trajectories of the Spike D614G mutation assuming different transmission fitness effects. Evolutionary trajectories are simulated under a two-strain SEIR epidemiological model. The epidemic is seeded with the 614D variant on January 15th while the 614G variant is allowed to enter the population through external introductions after February 15th. Because the rate at which 614D and 614G entered the US through external introductions largely determines the the rate at which the frequency of 614G grows, we explore different ratios of the external introduction rate for 614G relative to 614D.

**Supp. Figure 4.**
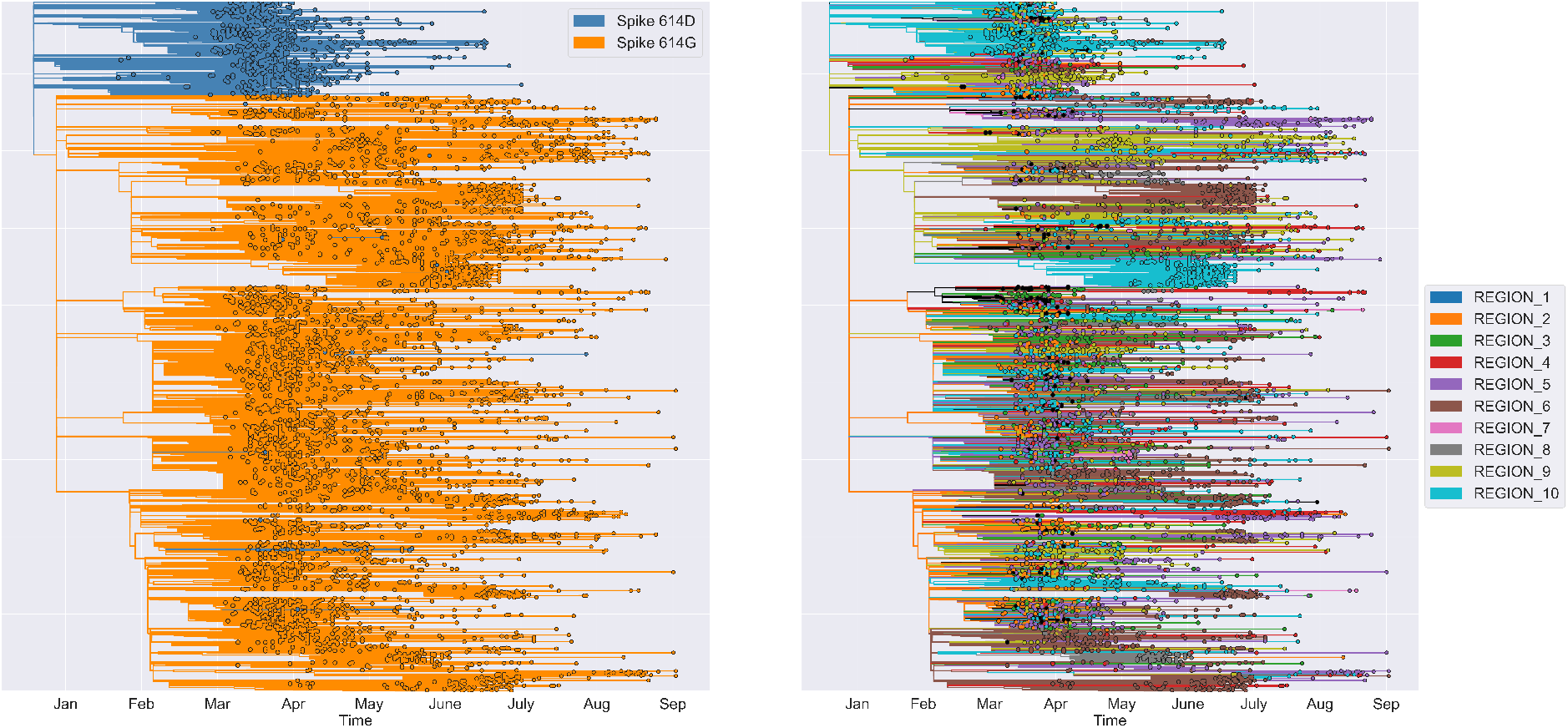
Maximum likelihood SARS-CoV-2 phylogenies with reconstructed ancestral features. The reconstructed Spike 614 variant type is shown on the left. The reconstructed geographic location in terms of US HHS Regions is shown on the left. The trees were thinned to only include 10% of all sampled tips in the full tree for purposes of visualization.

**Supp. Figure 5.**
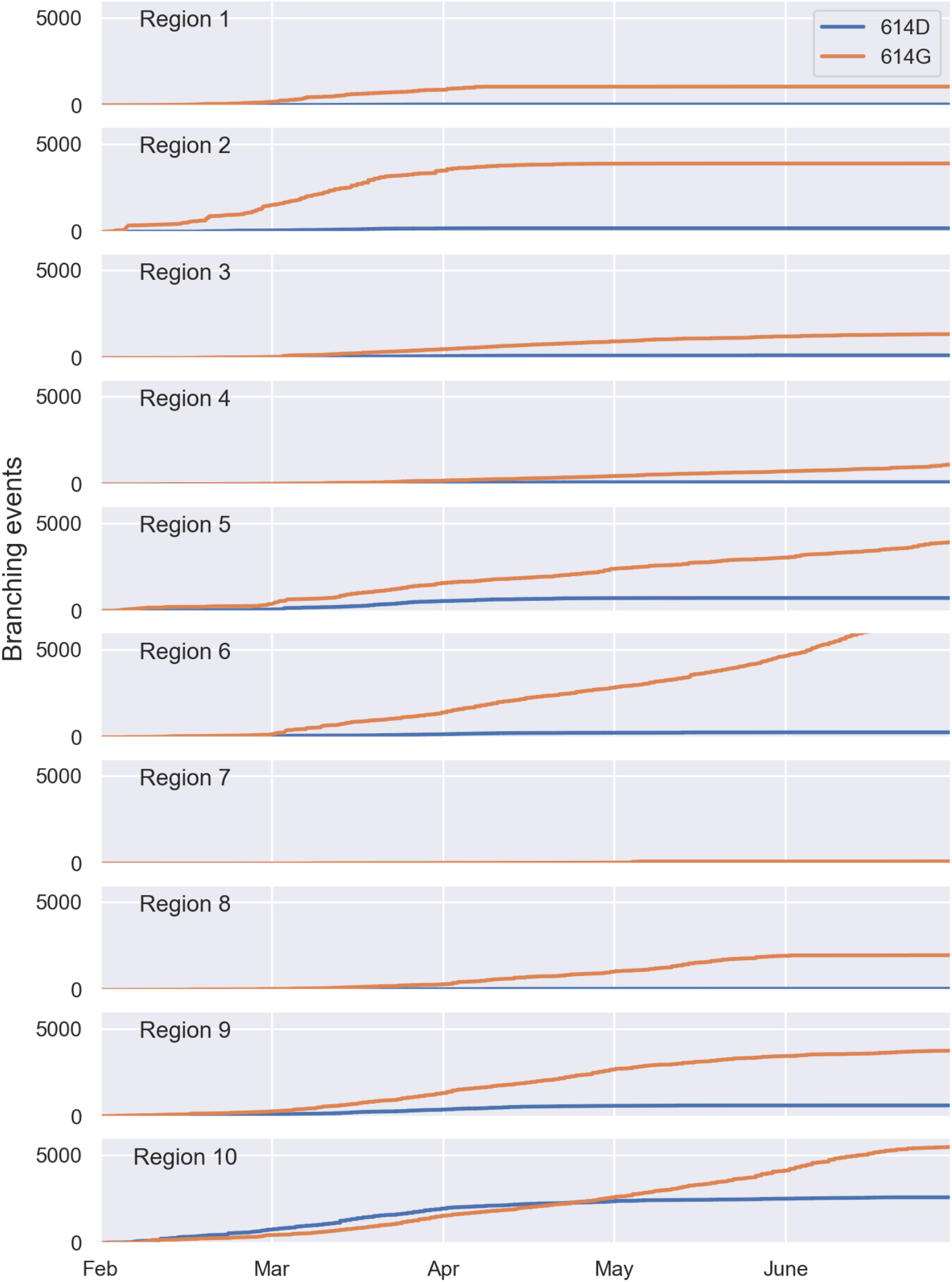
Cumulative branching events along lineages with the Spike 614 G versus D variant. Branching events are grouped by region based on their reconstructed ancestral location at the branching node.

**Supp. Figure 6.**
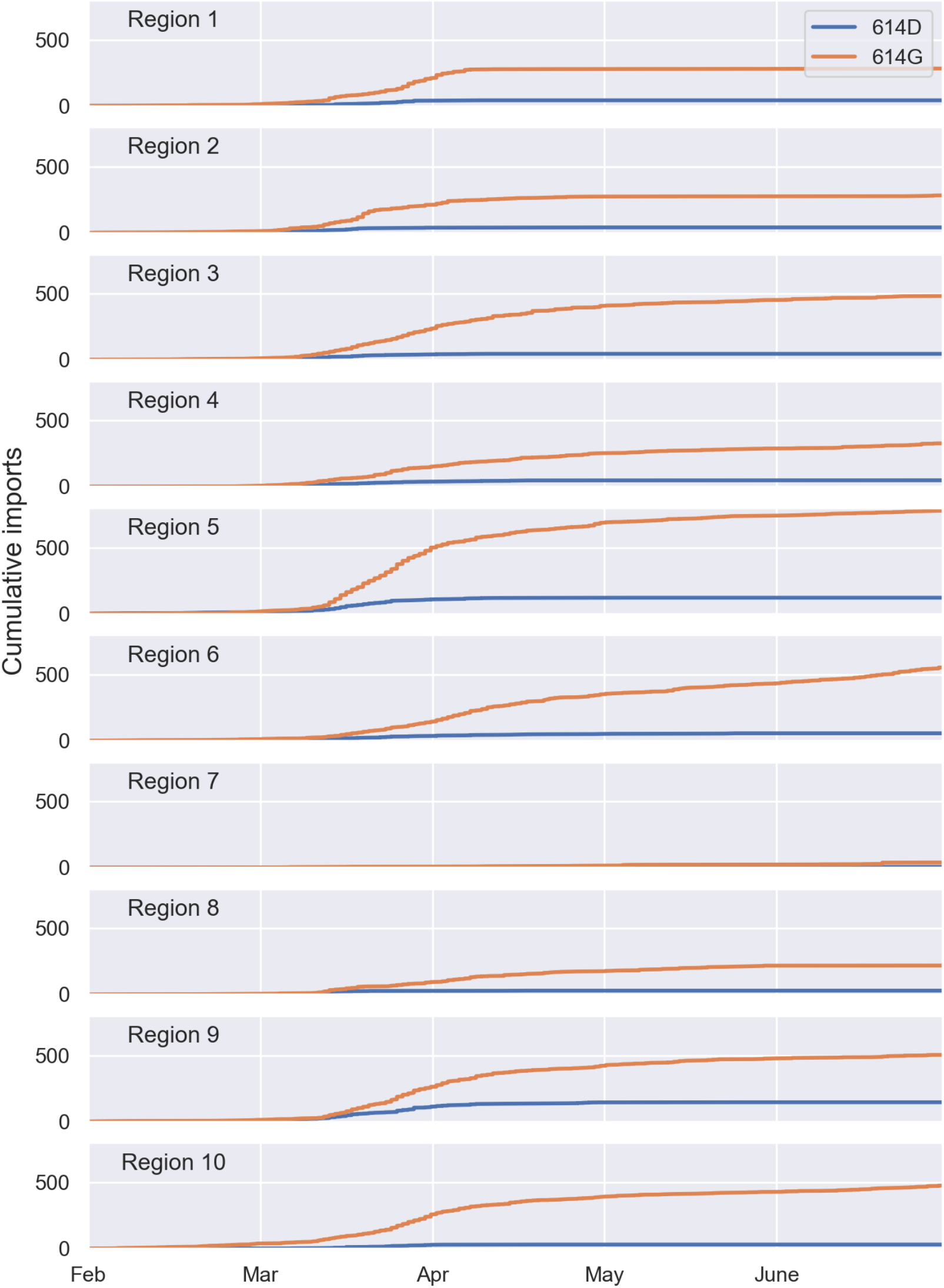
Cumulative number of lineages imported into each region with the Spike 614 G or D variant. Importation events were identified based on the reconstructed ancestral location of lineages in the phylogeny and defined here as a lineage’s ancestral location changing between a parent and a child node. The age/height of the child node was taken to be the time of the introduction event.

**Supp. Figure 7.**
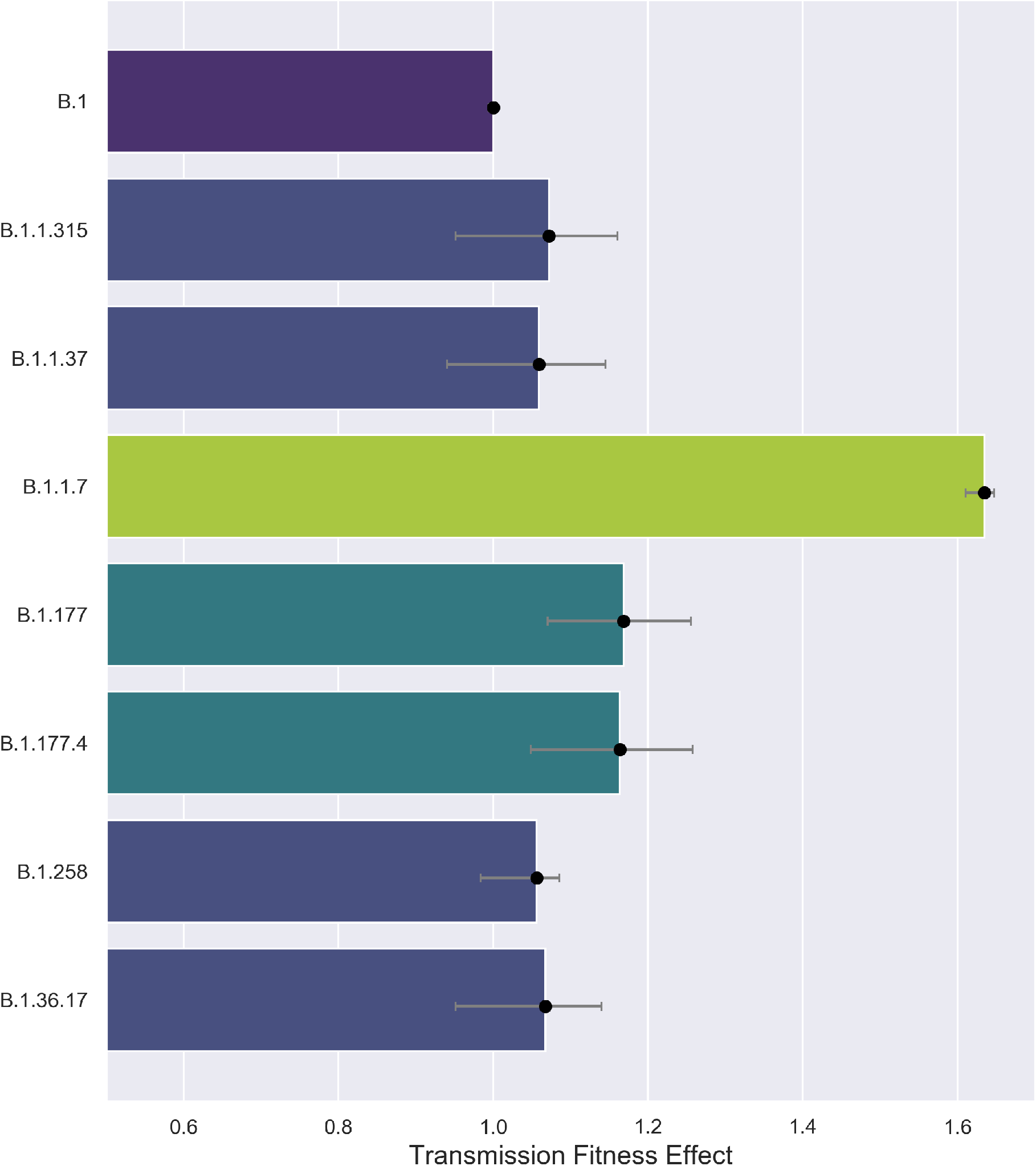
Estimated fitness of B.1.1.7 and other common lineages circulating in England. The fitness of each lineage is reported as a multiplicative effect on the base transmission rate shared among all lineages. Fitness effects are reported relative to B.1 in order to allow for equitable comparisons with the fitness of B.1.1.7 in the US.

**Supp. Figure 8.**
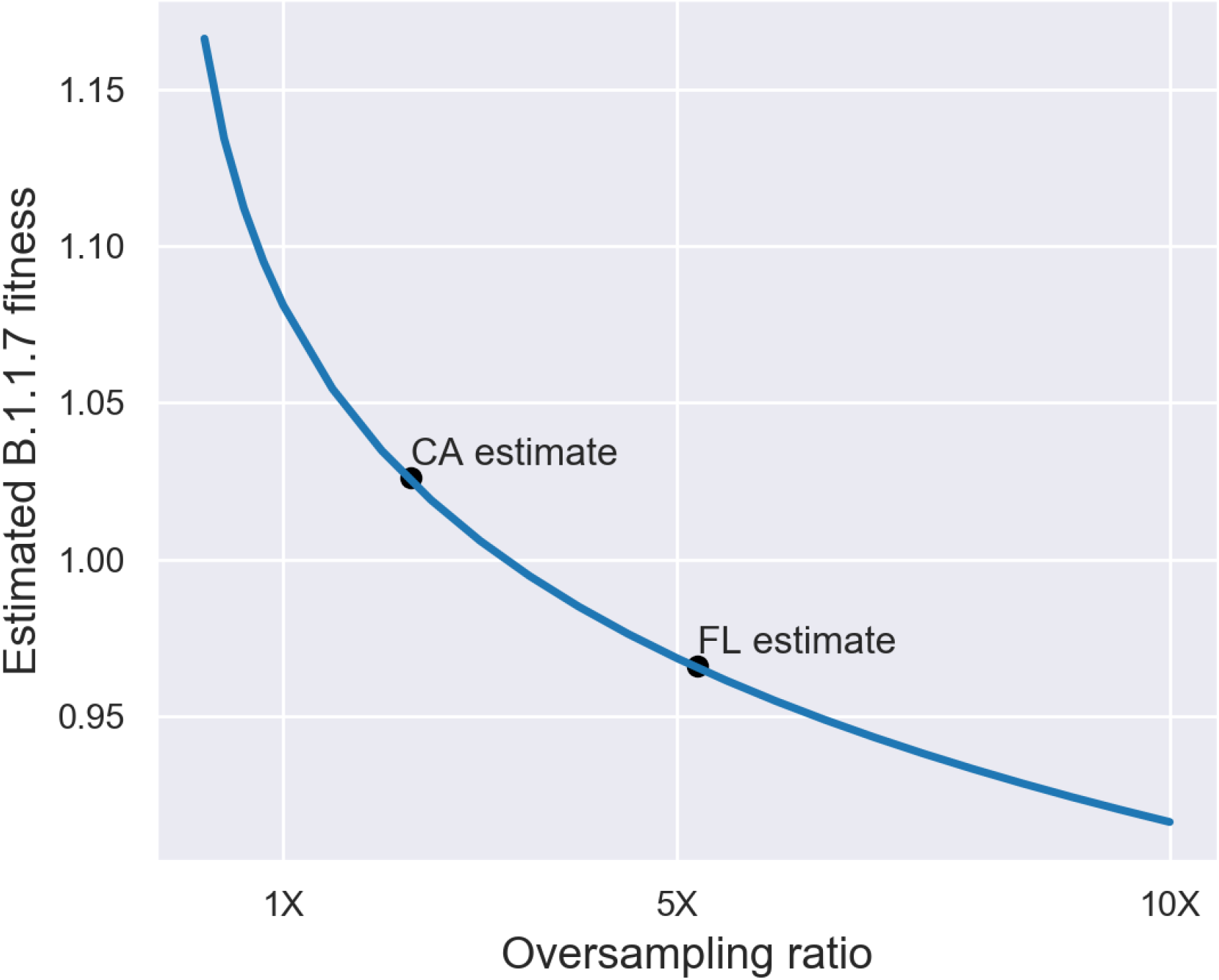
Relative fitness of B.1.1.7 estimated assuming different variant-specific sampling fractions. The oversampling ratio represents the sampling fraction of B.1.1.7 relative to all other sampled lineages. For comparison, the actual sampling fraction of B.1.1.7 was estimated for California (CA) and Florida (FL) based on the number of B.1.1.7 samples in GISAID relative to the total number of B.1.1.7 positive cases imputed using Helix’s qPCR/sequencing data.

**Supp. Figure 9.**
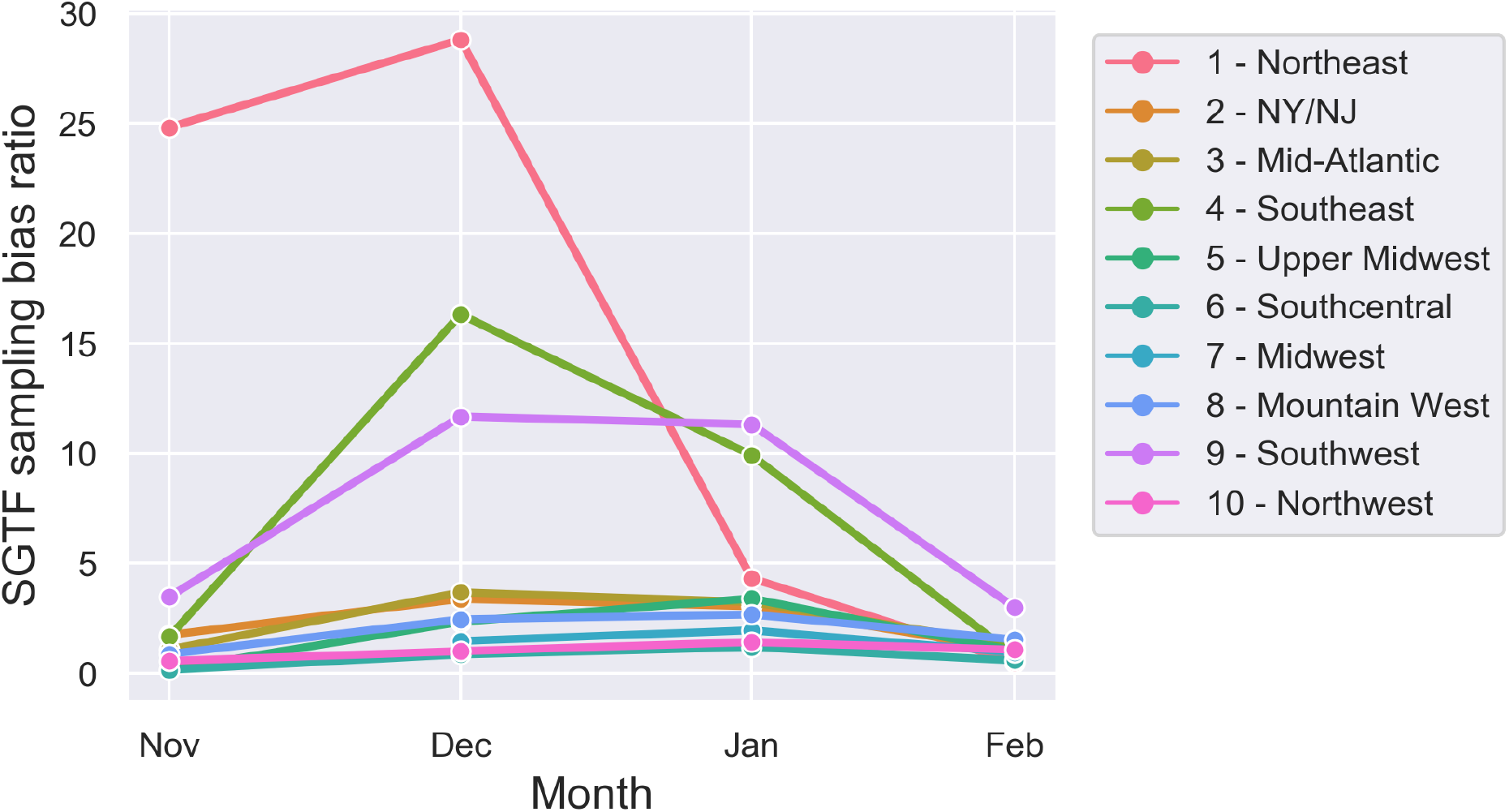
Sampling bias for variants with the Spike ΔH69/V70 mutation leading to SGTF. The SGTF sampling bias is computed as the ratio of the SGTF variant-specific sampling fraction over the non-SGTF variant sampling fraction for each region and time.

**Supp. Figure 10.**
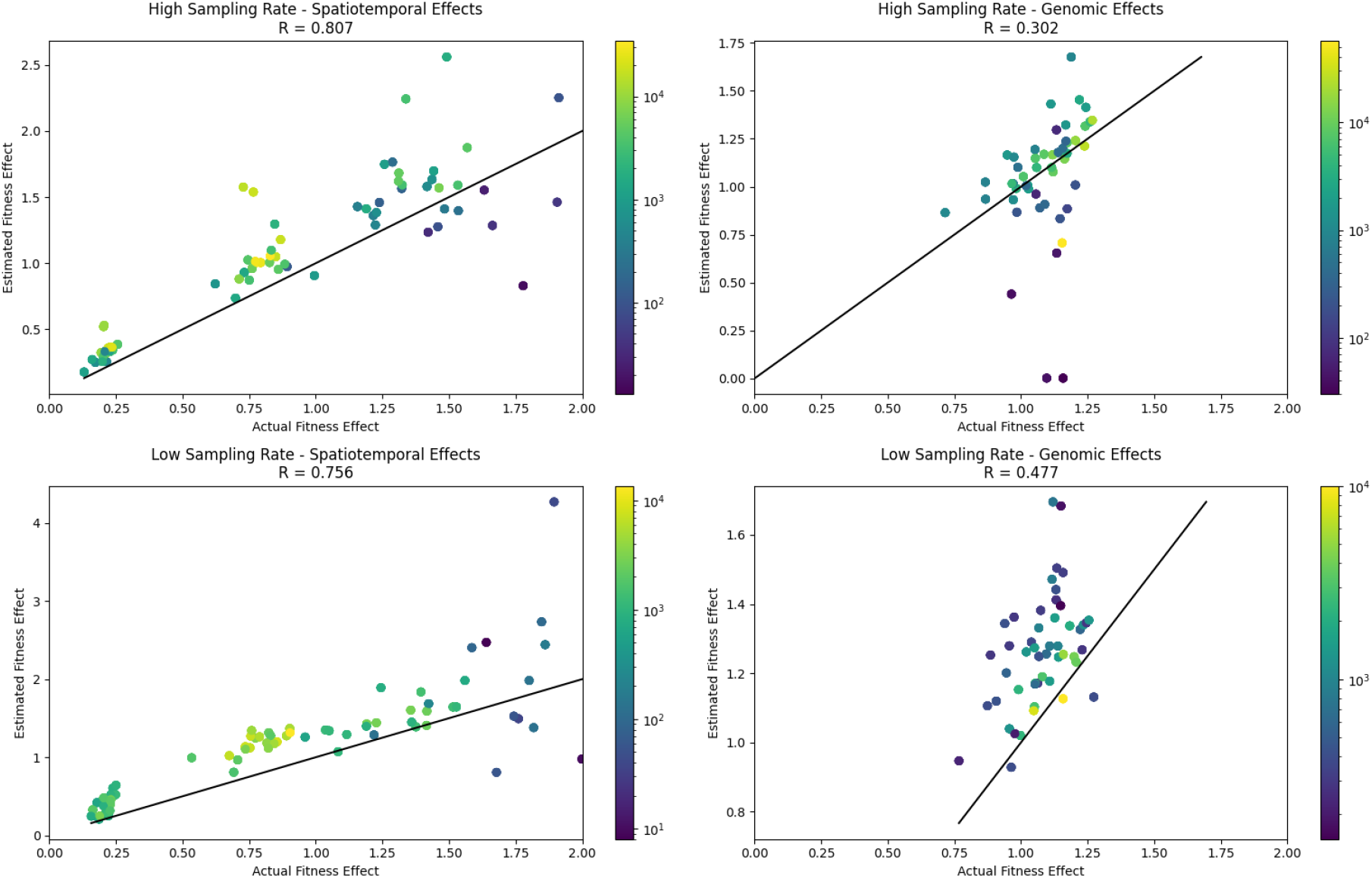
Actual (true) versus estimated fitness effects for features inferred from simulated phylogenetic trees. Dots represent the actual and estimated fitness value for an individual feature and are colored according to the number of individuals sampled with the corresponding feature. True fitness effects were known from simulations under a stochastic birth-death-sampling model with mutation. In these simulations, spatiotemporal (regional) and genomic (mutational) fitness effects were drawn randomly for each feature. In the top row sampling fractions are high (*σ* = *ρ* = 0.5) whereas in the bottom row sampling fractions are low (*σ* = *ρ* = 0.05). Simulation results were pooled across 10 simulations in each plot. Fitness estimates for features with a sampling fraction of less than 0.5% were discarded.

